# Flexible compression of sensory evidence makes human decisions more resilient to noisy neural computations

**DOI:** 10.64898/2026.10.27.762195

**Authors:** Julie Drevet, Lucas Benjamin, Lindsay Rondot, Jan Drugowitsch, Valentin Wyart

## Abstract

Categorizing ambiguous stimuli requires accumulating sensory evidence over time through noisy neural computations. Theories disagree on whether this evidence is accumulated in its native sensory space or in an abstract and compressed category space that retains only decision-relevant information. Here we designed a visual categorization task in which revealing decision categories before or after a stimulus sequence determines which space can be used to accumulate evidence, without changing the optimal decision. Humans were more accurate, and their inferences less noisy, when they could accumulate evidence in category space. Concurrent magnetoencephalographic (MEG) recordings revealed flexible and accuracy-predictive representations of evidence compressed onto decision categories in these conditions. Recurrent neural networks (RNNs) trained on the same task developed the same strategy, which made their decisions more resilient to noisy computations. Together, these findings suggest that the human brain mitigates the cost of noisy inferences by focusing limited computational resources on decision-relevant information.

## Introduction

Making accurate category-based decisions about ambiguous stimuli requires accumulating sensory evidence over time before committing to a decision. Evidence accumulation is widely seen as the core process that underlies perceptual decision-making under uncertainty^1^. Existing theories conceptualize this process as statistical inference^2,3^. To account for the pervasive suboptimality of human category-based decisions^4^, more recent theories introduce noise at different stages of the evidence accumulation process: noise in the sensory evidence being accumulated, but also in the accumulation (inference) process itself^5,6^. This second source of internal noise alone explains a surprisingly large fraction of human suboptimal decisions^7^.

But existing theories differ in terms of the representation space in which evidence is accumulated (**Fig. 1a**). Some theories propose that statistical inference is implemented by sensory cortices in the ‘native’ sensory space defined by sensory features – e.g., orientation for gratings or motion direction for random-dot patterns^8–10^. At the neural level, ‘bump attractor’^11–13^ and ‘population coding’^14^ models propose that sensory cortices can perform statistical inference and represent posterior distributions of sensory statistics in their activity – a proposal supported by studies decoding sensory uncertainty from human cortical activity^15–17^. By contrast, other theoretical accounts propose that evidence is accumulated in an ‘output’ (effector) space that encodes the action plan resulting from the decision^18,19^. Proponents of this view have reported that macaque monkeys postpone evidence accumulation when the mapping between choices and effectors is revealed only after stimulus offset^20^, although this finding does not appear to generalize to humans^21,22^.

**Figure 1.**
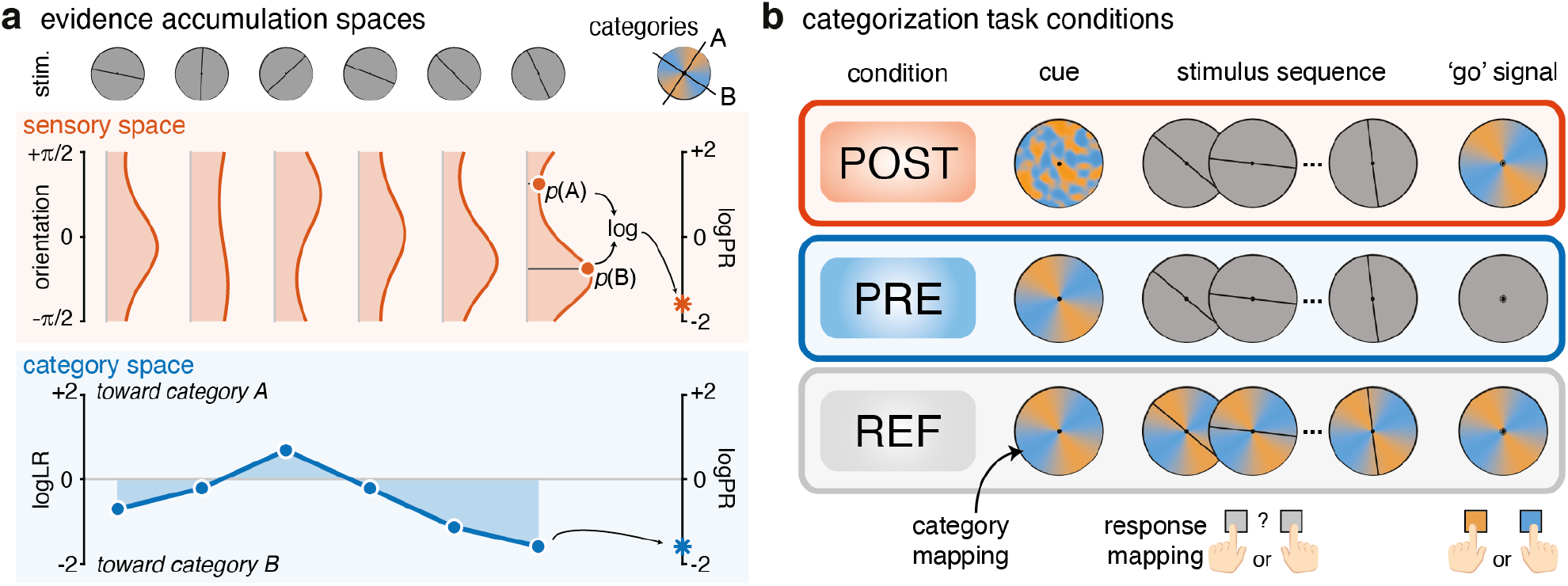
Representation of evidence during categorization. **(a)** Evidence accumulation spaces. A sequence of oriented bars (stimuli) is generated either by the category axis A or B. The optimal strategy to infer the underlying generative category of the sequence requires accumulating the information provided by each stimulus following Bayes’ rule. Accumulation in sensory space requires iteratively updating the full posterior distribution (which reduces to a two-dimensional vector of sufficient statistics, see Supplementary Information) over possible orientations for each presented stimulus. At the end of the sequence, the most likely category is computed from the posterior distribution using the log-posterior ratio log(*p*(*A*|sequence)⁄*p*(*B*|sequence)). Accumulation in category space implies projecting the information provided by each stimulus onto the A/B decision-relevant axis and updating the belief with this scalar quantity (logLR) only. At the end of the sequence, the most likely category is given by the sign of the log-posterior ratio (logPR) – here B – and is strictly equivalent in sensory and in category space. **(b)** Trial description. Participants had to categorize sequences of 4, 8 or 12 orientations drawn either from the orange or from the blue category presented at 3 Hz. The orientations of the orange and blue categories changed on every trial as indicated by a category-mapping cue presented either at the end (POST condition), before (PRE condition) or throughout (REF condition) the sequence. The response mapping was unknown during sequence presentation and changed at each trial. In POST, participants are forced to accumulate information in sensory space, but they can use category space in PRE and REF.

A third possibility is that evidence is accumulated neither in sensory nor effector space, but in an abstract and compressed ‘category’ space centered on the decision alternatives^23,24^. For two-alternative (A/B) decisions, this space corresponds to the geometrical projection of each piece of sensory evidence onto a decision axis that goes from category A on one end to category B on the other. This projection of sensory evidence onto the decision axis has an information cost, as it yields a compressed representation of sensory statistics that discards all dimensions irrelevant to the current decision. But it also has benefits when considering the cost of evidence accumulation, as only decision-relevant evidence needs to be accumulated. For A/B decisions, evidence accumulation in category space requires updating only a scalar log-posterior odds ratio rather than a full posterior distribution over feature values. If the neural resources available for evidence accumulation are limited, this reduction in dimensionality may allow devoting these resources to decision-relevant evidence – an idea reminiscent of efficient coding principles in sensory systems^25^. Accumulating evidence in category space also requires flexibility, since sensory evidence must be projected onto a new decision axis whenever the decision categories change. To our knowledge, no study has compared conditions that explicitly manipulate the possibility of accumulating evidence in these different representation spaces. It therefore remains unclear whether humans prefer one of these spaces, and whether this preferred space yields more accurate decisions for the exact same amount of sensory evidence.

To address these questions, we devised a visual categorization task in which the possibility of accumulating evidence in sensory or category space can be experimentally manipulated without changing the optimal decision (**Fig. 1b**). By providing decision categories either after or before stimulus presentation, we could compare the behavioral, computational and MEG signatures of evidence accumulation performed in sensory or category space. Our findings show that humans – and recurrent neural networks (RNNs) trained in the same conditions – preferentially accumulate sensory evidence in category space. This compression is associated with more precise inferences in humans, and makes RNNs more resilient to noisy recurrent computations.

## Results

### Visual categorization task

To examine how sensory evidence is represented during category-based decisions, we designed a visual categorization task that manipulates whether participants can represent it in its native sensory space or in a compressed category space. Participants were asked to categorize sequences of oriented stimuli (**Fig. 1b**). Each orientation sequence was drawn from one of two overlapping probability distributions centered on orthogonal mean orientations, each associated with a color-coded decision category (A: orange, B: blue; **Supplementary Fig. 1**). The mean orientations of the two decision categories changed randomly from one sequence to the next, and were indicated by a category-mapping cue (a color wheel). Accurate decisions therefore required participants to flexibly relate sensory evidence to a new decision axis on every trial. Participants had to accumulate the sensory evidence provided by each oriented stimulus, and report the category of the sequence when prompted by a ‘go’ signal after sequence offset (see Methods). Because the mapping between categories and response buttons was randomized across trials and revealed only on the ‘go’ signal, evidence could not be accumulated in the effector space in any condition.

In the first condition (POST; **Fig. 1b**), decision categories were revealed only with the ‘go’ signal after the offset of the sequence, constraining evidence accumulation to sensory space – i.e., computing the posterior distribution over the mean orientation of the sequence until the decision categories are revealed. In the second condition (PRE; **Fig. 1b**), decision categories were briefly presented before sequence onset, allowing evidence accumulation in category space – i.e., projecting the sensory evidence provided by each stimulus onto the decision axis defined by the two categories, whose mean orientations must be maintained in working memory. In the third, control condition (REF; **Fig. 1b**), decision categories were displayed throughout the trial, as a color wheel in the background of the oriented stimuli until the response.

For a given orientation sequence, the Bayes-optimal strategy yields the same decision, with the same uncertainty, in all three conditions. Indeed, the POST condition requires computing the posterior distribution over the mean orientation of the sequence (evidence accumulation in sensory space) and then comparing the posterior probabilities of the two decision categories revealed with the ‘go’ signal. This is mathematically equivalent to projecting each stimulus onto the decision axis and accumulating the resulting category evidence across the sequence (evidence accumulation in category space; **Fig. 1a**; see **Supplementary Information**) – a strategy possible only in the PRE and REF conditions. However, whereas accumulating evidence in sensory space requires updating a full probability distribution after each stimulus (which reduces to a two-dimensional vector of sufficient statistics, see Supplementary Information), accumulating evidence in category space requires updating only a scalar log-posterior odds ratio. In other words, although normatively equivalent, accumulating evidence in these two representation spaces relies on representations of different dimensionality, which may be differentially prone to suboptimalities such as inference noise^5,7^.

### Behavioral signatures of evidence accumulation in category space

We reasoned that if participants accumulate evidence in sensory space in all three conditions, their decision accuracy should be higher in POST and REF, where decision categories are displayed at the ‘go’ signal, than in PRE, where they must be maintained in working memory. By contrast, if participants preferentially accumulate evidence in category space, their decision accuracy should be higher in PRE than in POST, where this is impossible, and even higher in REF, where it is not necessary to maintain decision categories in working memory. Accuracy was suboptimal in all three conditions (Bayes-optimal accuracy: 83.5%), with a pattern consistent with a preference for evidence accumulation in category space (POST: 70.9 ± 0.7%; PRE: 75.2 ± 0.7%; REF: 78.4 ± 0.5%; mean ± s.e.m.; **Fig. 2a**). Indeed, decision accuracy differed significantly between conditions (repeated-measures ANOVA: F_2,72_ = 58.2, *p* < 0.001, partial *η*^2^ = 0.618), and was significantly higher in PRE than in POST (post-hoc paired *t*-test: *t*_36_ = 6.2, *p* < 0.001, Cohen’s *d* = 1.02, 95% CI = [2.62% 6.00%]). As expected, accuracy was even higher in REF, where decision categories are displayed throughout the sequence and need not be held in working memory (*t*_36_ = 4.74, *p* < 0.001, Cohen’s *d* = 0.77, 95% CI = [1.59% 4.96%]). Analyses of response times further supported these conclusions (see **Supplementary Results** and **Supplementary Fig. 2**). In particular, the decision-irrelevant evidence provided by the sequence – computed relative to the boundaries between the two categories – slowed down responses in POST, but not in PRE and REF (**Supplementary Fig. 2d**).

**Figure 2.**
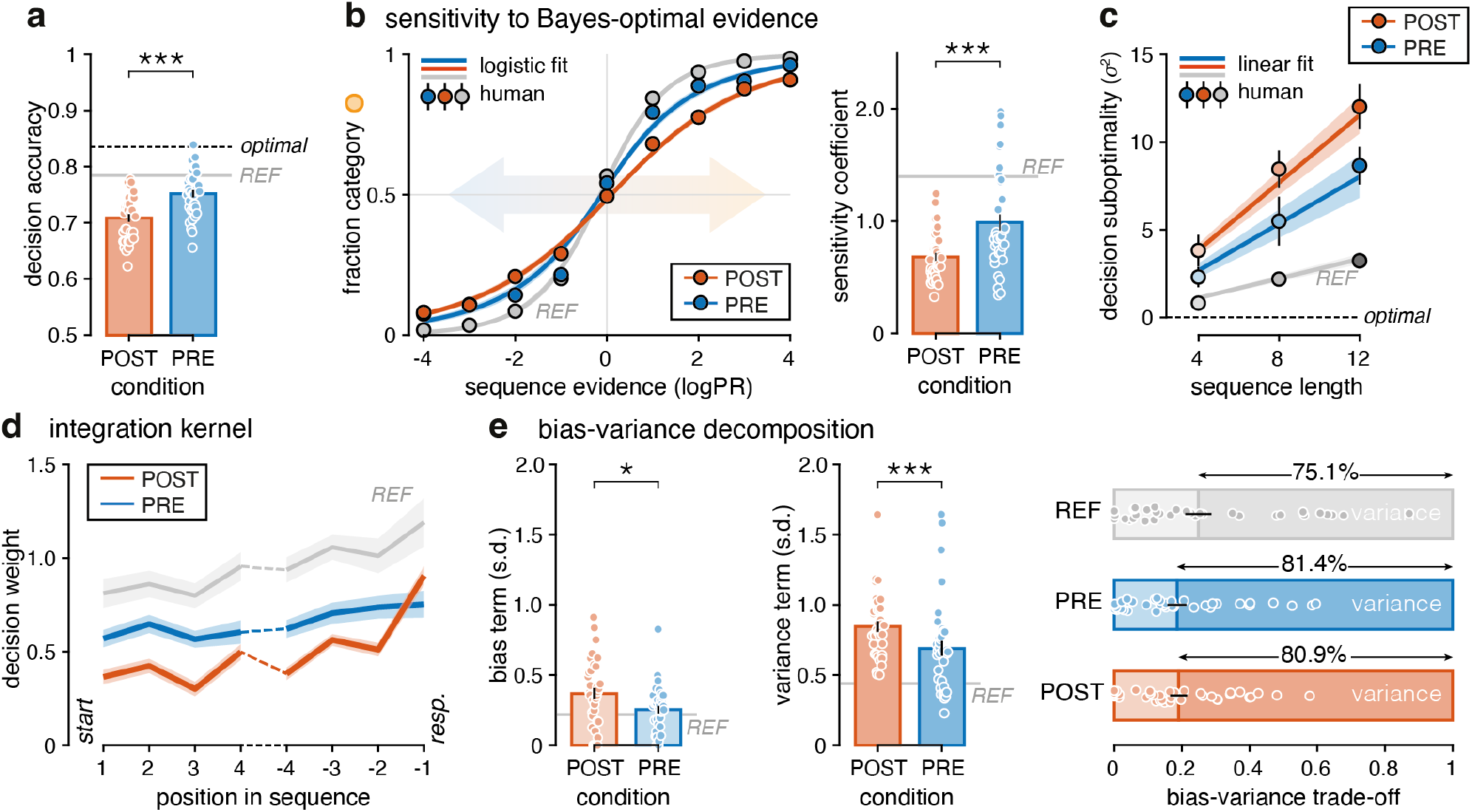
Effects of accumulation space on human inference (N = 37). **(a)** Decision accuracy. Participants are suboptimal in all conditions, with lowest performance in POST and best performance in REF. **(b)** Sensitivity to Bayes-optimal evidence. Left: Fraction of responses in favor of the orange category as a function of whole-sequence evidence. Positive log-posterior ratios (logPR) are in favor of the orange category. Color lines indicate best-fitting logistic regression. Right: Sensitivity coefficient (slope of the logistic regression). Sensitivity is lowest in POST. **(c)** Inference-driven source of decision suboptimality. Estimated decision suboptimality (dots and bars: mean ± s.e.m.) grows with sequence length in all conditions. Suboptimality is larger and grows faster in POST. Lines indicate model predictions. **(d)** Integration kernel. Evidence weight as a function of sample position from sequence start and from response. Late samples contribute more than early samples to participants’ decision in all conditions. **(e)** Bias-variance decomposition of decision suboptimality. Left: Bias term – deterministic deviations – is higher in POST. Middle: Variance term – random deviations – is higher in POST. Right: Relative proportion of variability due to deterministic (light colors) and random (full colors) deviations. In all panels: the dashed black lines correspond to an optimal inference process, the solid gray lines correspond to participants’ behavior in REF, large dots, bars and error bars indicate mean ± s.e.m., small dots indicate single participants, shaded areas correspond to s.e.m., three stars and one star indicate a significant effect at *p* < 0.001 and *p* < 0.05, respectively.

To understand why participants were less accurate in POST, where evidence can only be accumulated in sensory space, we measured the sensitivity of their decisions to the category evidence provided by each sequence (i.e., the total log-posterior odds ratio). Decisions were less sensitive to this evidence in POST than in PRE (*t*_36_ = −5.37, *p* < 0.001, Cohen’s *d* = −0.88, 95% CI = [−0.46 −0.17]; **Fig. 2b**). In all conditions, decision suboptimality – the estimated variance of deviations from Bayes-optimal decisions – increased with the number of stimuli in the sequence (F_2,72_ = 56.1, *p* < 0.001, partial *η*^2^ = 0.609; **Fig. 2c**) – a diagnostic signature of noise arising during evidence accumulation rather than during response selection^5,7^. Importantly, this scaling was steeper in POST (F_4,144_ = 4.06, *p* = 0.011, partial *η*^2^ = 0.101), indicating a larger suboptimality of evidence accumulation in this condition. This suboptimality can arise from random variability, but also from systematic biases such as uneven weighting of evidence samples over time^7,13,26–28^. To quantify these temporal biases, we estimated the decision weight given to each evidence sample depending on its position in the sequence. We found a moderate recency effect (F_7,252_ = 15.38, *p* < 0.001, partial *η*^2^ = 0.299; **Fig. 2d**), which was more pronounced in POST (F_14,504_ = 3.07, *p* = 0.0065, partial *η*^2^ = 0.078). This suggests that accumulated evidence decays more quickly when it is accumulated in sensory space.

Other unidentified biases may also contribute to the suboptimality of evidence accumulation. To decompose decision suboptimality into bias and variance components in each condition, we used a validated method^5,7^ based on the consistency of participants’ decisions across two presentations of the exact same stimulus sequences (see **Methods**). This method provides estimates of the magnitude of the bias and variance components in each condition, both of which were larger in POST than in PRE (and REF) conditions (bias: F_2,72_ = 8.2, *p* = 0.0023, partial *η*^2^ = 0.187; variance: F_2,72_ = 48.0, *p* < 0.001, partial *η*^2^ = 0.571; **Fig. 2e**). Thus, accumulating evidence in category space – in PRE and REF – is associated with reductions of both the bias and the variance components of decision suboptimality. In all conditions, the variance component explained most of participants’ decision suboptimality (POST: 80.9 ± 2.5%; PRE: 81.4 ± 2.8%; REF: 75.1 ± 3.8%).

### Computational modeling of suboptimal evidence accumulation

To explain these behavioral observations, we modeled evidence accumulation as a suboptimal inference process^7,29–31^, with three optional parameters (**Fig. 3a**): 1. inference imbalance *γ*, capturing recency (or primacy) effects through a deterministic imbalance in the weighting of the prior belief at each inference step (i.e., after each stimulus), 2. inference noise *σ*_inf_, capturing random variability at each inference step, and 3. selection noise *σ*_sel_, capturing random variability at response selection (see Methods). Because accumulating evidence in sensory and category spaces is mathematically equivalent in our task, each parameter has the same impact on decisions in both spaces and can therefore be compared across conditions (see **Supplementary Methods** and **Supplementary Fig. 3**).

**Figure 3.**
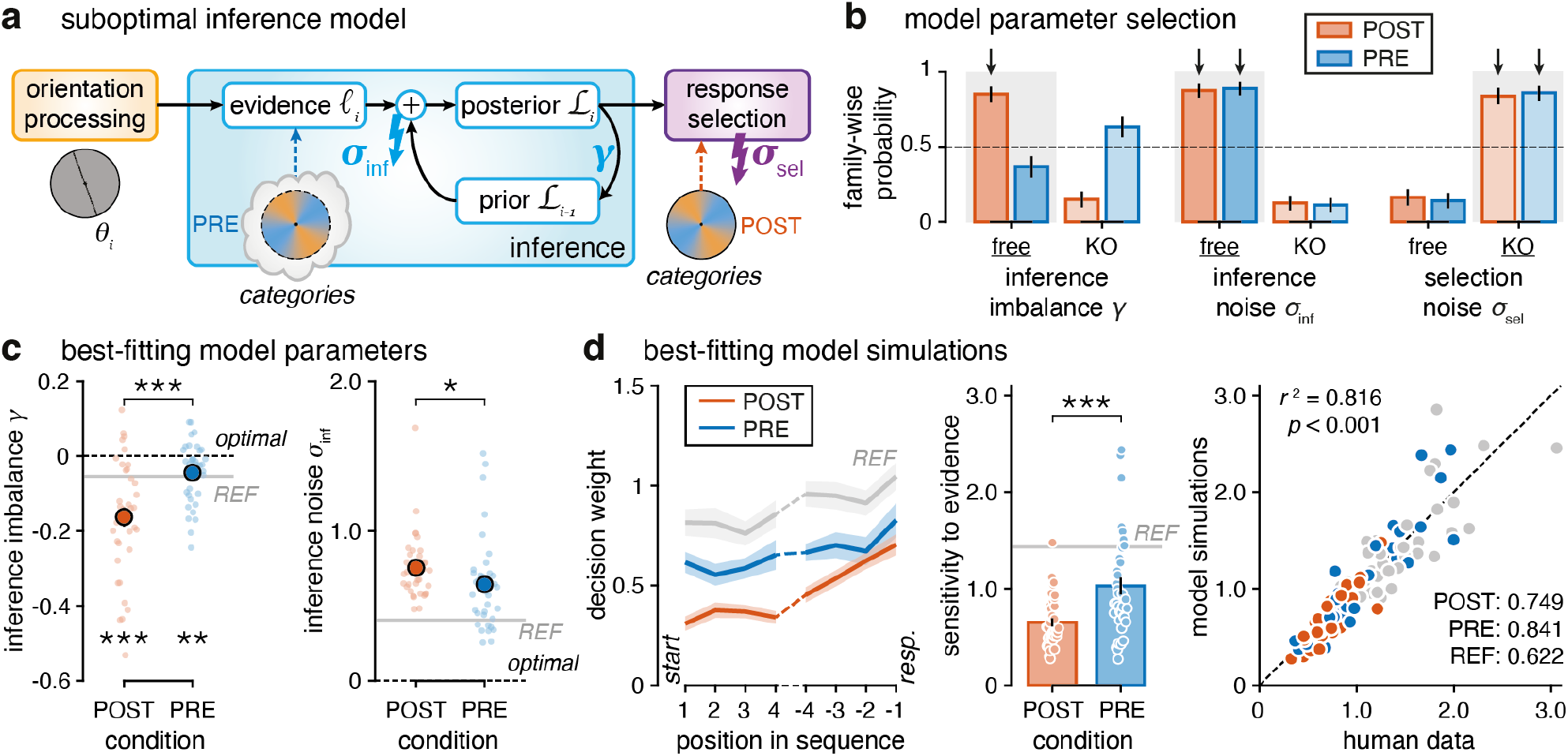
Modeling suboptimal cognitive inference. **(a)** Suboptimal inference model. Optimal Bayesian inference estimates the log-likelihood ℓ_*i*_ provided by each orientation *θ*_*i*_, accumulates this evidence across the *n* stimuli of the sequence to obtain a log-posterior *ℒ*_*n*_, and selects the most likely category. Suboptimality is introduced through three parameters: 1. the inference imbalance *γ* weighting the log-prior *ℒ*_*i−1*_ before being combined with ℓ_*i*_; 2. the inference noise *σ*_inf_ added to each accumulated ℓ_*i*_; and 3. the selection noise *σ*_sel_ added to the final log-posterior ℒ_*n*_. In PRE, the categories can be used to compute a compressed format for the evidence. In POST, the categories are provided at the end of the inference process. The log-prior of the first stimulus is *ℒ*_0_ = 0. **(b)** Factorized Bayesian model selection. Family-wise probabilities of the model variants in which each parameter is freely fitted or knocked out (KO, fixed to 0). Inference imbalance improves fit in POST but not in PRE, inference noise explains participants’ suboptimal choices, selection noise does not. Bars show means ± s.d. of estimated Dirichlet distributions, black arrows indicate exceedance probabilities above 99%. Gray shaded areas indicate selected model parameters. **(c)** Best-fitting model parameters. Left: Participants’ inference is leaky (*γ* < 0 in all conditions: POST: −0.163 ± 0.025; PRE: −0.045 ± 0.013; REF: −0.055 ± 0.013, mean ± s.e.m.). Right: Participants’ inference is noisy (*σ*_inf_: POST: 0.755 ± 0.036; PRE: 0.644 ± 0.051; REF: 0.406 ± 0.014, mean ± s.e.m.). Large color dots with error bars (hidden behind) indicate mean ± s.e.m., dashed black lines correspond to an optimal inference process. **(d)** Best-fitting model simulations reproducing humans’ behavior. Left: Simulated integration kernel (for humans, see Fig. 2d). Middle: Sensitivity to evidence (for humans, see Fig. 2b). Right: Correlation between simulated and humans’ sensitivity coefficients in each condition, with condition-specific *r*^2^. In (c,d): the gray lines and dots correspond to REF, small color dots indicate single participants, error bars and shaded areas indicate s.e.m., three stars indicate a significant effect at *p* < 0.001, two stars at *p* < 0.01 and one star at *p* < 0.05.

After verifying that each parameter could be recovered from synthetic behavior^7,32,33^ (see **Methods** and **Supplementary Fig. 4a**), we used factorized Bayesian model selection to assess which of the three sources of suboptimality best explained participants’ behavior (**Fig. 3b**). This procedure confirmed the presence of inference noise and ruled out selection noise in all conditions (model variants with *σ*_inf_ fitted to behavior: exceedance *p* > 0.999; model variants with *σ*_sel_ fixed to 0: exceedance *p* > 0.999). The inference imbalance parameter improved fits in POST (model variants with *γ* fitted to behavior: exceedance *p* > 0.999), but not in PRE (exceedance *p* = 0.034) where this parameter had an estimated prevalence of 36.7%. For further analyses, we selected the model variant with inference imbalance *γ* and inference noise *σ*_inf_, but without selection noise *σ*_sel_.

We used this model variant in all conditions to compare parameter estimates across conditions. First, we found that in all conditions, most participants underweighted previously accumulated evidence, consistent with the recency effect described above (*γ*: *t*-test to 0: *t*_36_ < −3.3 in all conditions, PRE: *p* < 0.01, POST and REF: *p* < 0.001; **Fig. 3c**, left). The inference imbalance *γ* differed across conditions (F_2,72_ = 17.5, *p* < 0.001, partial *η*^2^ = 0.327), with stronger underweighting of accumulated evidence in POST, as expected from the bias-variance decomposition (POST-PRE post-hoc *t*-test: *t*_36_ = −4.5, *p* < 0.001, Cohen’s *d* = −0.74, 95% CI = [−0.182 −0.054]) and similar imbalance in PRE and REF, where the categories were shown before accumulation (PRE-REF post-hoc *t*-test: *t*_36_ = 1.03, *p* = 0.55, Cohen’s *d* = 0.17, 95% CI = [−0.014 +0.035], BF_01_ = 3.43), suggesting that maintaining the category axis in working memory does not affect the amount of bias in evidence accumulation. Second, inference noise corrupted evidence accumulation in all conditions (**Fig. 3c**, right), even when accounting for the inference imbalance (**Supplementary Fig. 4c**). Inference noise also differed across conditions (F_2,72_ = 35.96, *p* < 0.001, partial *η*^2^ = 0.499). Importantly, inference was less precise when less information about the generative categories was available during accumulation (POST-PRE: *t*_36_ = 2.53, *p* = 0.042, Cohen’s *d* = 0.41, 95% CI = [0.003 0.217], BF_10_ = 2.83; PRE-REF: *t*_36_ = 5.07, *p* < 0.001, Cohen’s *d* = 0.83, 95% CI = [0.12 0.36], BF_10_ = 1752). Unlike the imbalance, inference noise increased when the category axis had to be maintained in working memory, suggesting that inference noise in PRE includes noise arising from the imprecise maintenance of the category axis in working memory. Simulations of the best-fitting model reproduced participants’ behavioral signatures (**Fig. 3d** and **Supplementary Fig. 5**). These findings corroborate the idea that accumulating evidence in category space is more precise than accumulating the same evidence in sensory space.

### Neural signatures of evidence accumulation in category space

We next examined the neural signatures of stimulus processing in sensory space (POST) and category space (PRE) using magnetoencephalography (MEG). To do so, we trained ridge regression models to decode different characteristics of each stimulus from stimulus-locked spatial patterns of MEG activity (see **Methods**). We focused statistical comparisons on POST and PRE, which are matched in visual stimulation during the sequence. REF, in which the category-mapping cue was displayed behind each stimulus, is shown for reference. We could decode stimulus orientation from early MEG responses, peaking 90 ms after stimulus onset (**Fig. 4a**), with no significant difference in peak coding precision between POST and PRE (jackknifed *t*_36_ = 1.46, *p* = 0.15, BF_01_ = 2.13). Using structural brain scans of each participant, we localized the brain regions supporting this decoding to V1 and surrounding early visual cortex (**Fig. 4a**; see **Methods**). This similar decoding of stimulus orientation in POST and PRE suggests that early visual processing does not differ between the two conditions.

**Figure 4.**
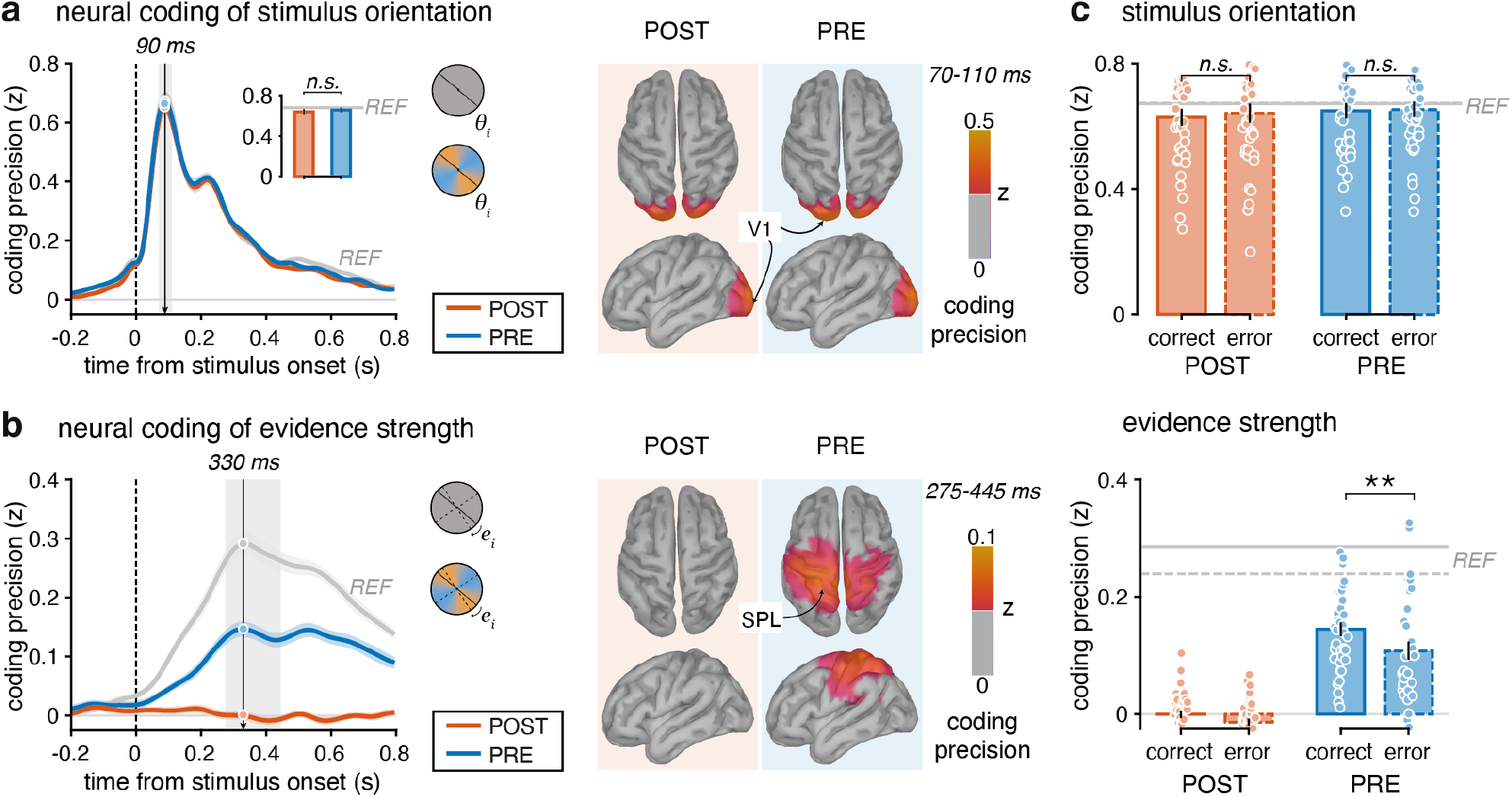
Neural correlates of stimulus processing (N = 37). **(a)** Neural coding of stimulus orientation. Left: Time course of coding from whole-brain MEG signals. Stimulus orientation coding peaks around +90 ms (jackknifed mean: POST: 88.6 ms; PRE: 88.4 ms; REF: 89.0 ms; jackknife-corrected F_2,72_ = 0.44, *p* = 0.63) with similar precision between PRE and POST. The gray shaded area indicates the [+70 +110] ms time window, during which the coding precision reaches 90% of its peak value in each condition. Top right inset: Coding averaged on this time window. No difference found between PRE and POST. Middle: Searchlight-based coding across the cortical surface (thresholded at 50% of the PRE coding peak), peaking in occipital cortex surrounding V1. **(b)** Neural coding of evidence strength. Same conventions as in (a). Left: Evidence strength peaks around +330 ms in PRE and REF (jackknifed mean: PRE: 326 ms; REF: 332 ms) but is not decoded in POST. The coding precision reaches 90% of its peak value in at least one condition on [+275 +445] ms (condition-specific: PRE: [+275 +395] ms (first peak), REF: [+280 +445] ms). Middle: Coding peaks in parietal cortex surrounding the SPL in PRE and is absent in POST. **(c)** Comparison of coding precision between correct and error trials. Coding is averaged on the time windows defined by the gray shaded area in (a,b). Evidence strength (bottom), but not stimulus orientation (top), is coded more precisely before correct than error decisions in PRE and REF. In all panels: solid lines and shaded areas, bars and error bars indicate mean ± s.e.m., small dots indicate single participants, the gray lines correspond to REF, two stars indicate a significant effect at *p* < 0.01 and n.s. indicates no statistical difference for one-tailed *t*-test testing correct > error.

We then decoded the strength of the evidence provided by each stimulus in favor of either category – a quantity that can be computed only when categories are known during evidence accumulation. As expected, we could not decode evidence strength from MEG activity in POST. By contrast, we could decode evidence strength in PRE from later MEG responses, peaking 330 ms after stimulus onset (cluster-corrected *p* < 0.001; **Fig. 4b**). Unlike stimulus orientation, evidence strength was best decoded from parietal cortex, centered on the superior parietal lobule (SPL) and the intraparietal sulcus (IPS). This finding is consistent with the idea that humans accumulate evidence in category space when the categories are known. Because the category axis changed from trial to trial, decoding evidence strength across trials relied on a neural code shared across stimulus-category mappings – i.e., abstracted from the specific orientations that define the categories on each trial.

We then tested whether the neural coding of these two stimulus characteristics during evidence accumulation predicted the accuracy of the resulting decision at the end of the trial (**Fig. 4c** and **Supplementary Fig. 6**). The decoding of stimulus orientation was not stronger before correct decisions than before errors in either condition (POST: one-tailed *t*_36_ = −1.9, *p* = 0.96; PRE: one-tailed *t*_36_ = −0.5, *p* = 0.71). By contrast, the decoding of evidence strength in PRE was stronger before correct decisions than before errors (one-tailed *t*_36_ = 3.31, *p* = 0.0011). This brain-behavior association is consistent with the idea that the neural representation of evidence in category space supports accurate category-based decisions.

### Neural compression of accumulated evidence in category space

The two characteristics decoded above are computed upstream from the inference process. In this task, inference consists of updating the belief about the generative statistics of the stimulus sequence with the evidence provided by each new stimulus. To examine the neural computation of belief updates in sensory and category spaces, we decoded the direction of belief updates from MEG activity – ranging from positive updates (when the incoming evidence is consistent with the current belief) to negative updates (when the incoming evidence is conflicting with the current belief). Because belief updates can be defined with respect to any pair of categories, we computed them separately for the ‘true’ (decision-relevant) categories, and for ‘null’ (decision-irrelevant) categories rotated by 45° (see **Methods**). Because the null categories are centered on the boundaries between the true categories, the evidence provided by each stimulus relative to the null categories corresponds to the component of its orientation that is orthogonal to the decision axis, and thus irrelevant to the decision. In POST, where the categories are unknown during the sequence, differences between the coding of belief updates relative to the true and null categories cannot reflect category-specific processing, and thus provide a baseline for PRE.

We could decode belief updates relative to the true categories more precisely in PRE than in POST, i.e., more precisely when evidence could be accumulated in category space (cluster-corrected *p* < 0.001; **Fig. 5a**), peaking 210 ms after stimulus onset. In POST, belief updates relative to the true categories could also be decoded, but not more precisely than belief updates relative to the null categories – consistent with the idea that MEG activity in POST reflects the consistency of each stimulus with the evidence accumulated so far in sensory space, which does not depend on the categories. Their decoding was distributed across the brain, including the parietal regions coding for evidence strength and extending to temporal and prefrontal regions (**Supplementary Fig. 7a**). By contrast, we could decode belief updates relative to the null categories less precisely in PRE than in POST, i.e., less precisely when evidence could be accumulated in category space (cluster-corrected *p* < 0.01; **Fig. 5a**). These two opposing effects indicate that evidence accumulation in category space is associated with a neural ‘compression’ of accumulated evidence, i.e., enhanced coding of belief updates along the decision-relevant axis and reduced coding along the decision-irrelevant axis. This compression does not imply that sensory information is discarded at early processing stages: the early coding of stimulus orientation did not differ between PRE and POST (**Fig. 4a**).

**Figure 5.**
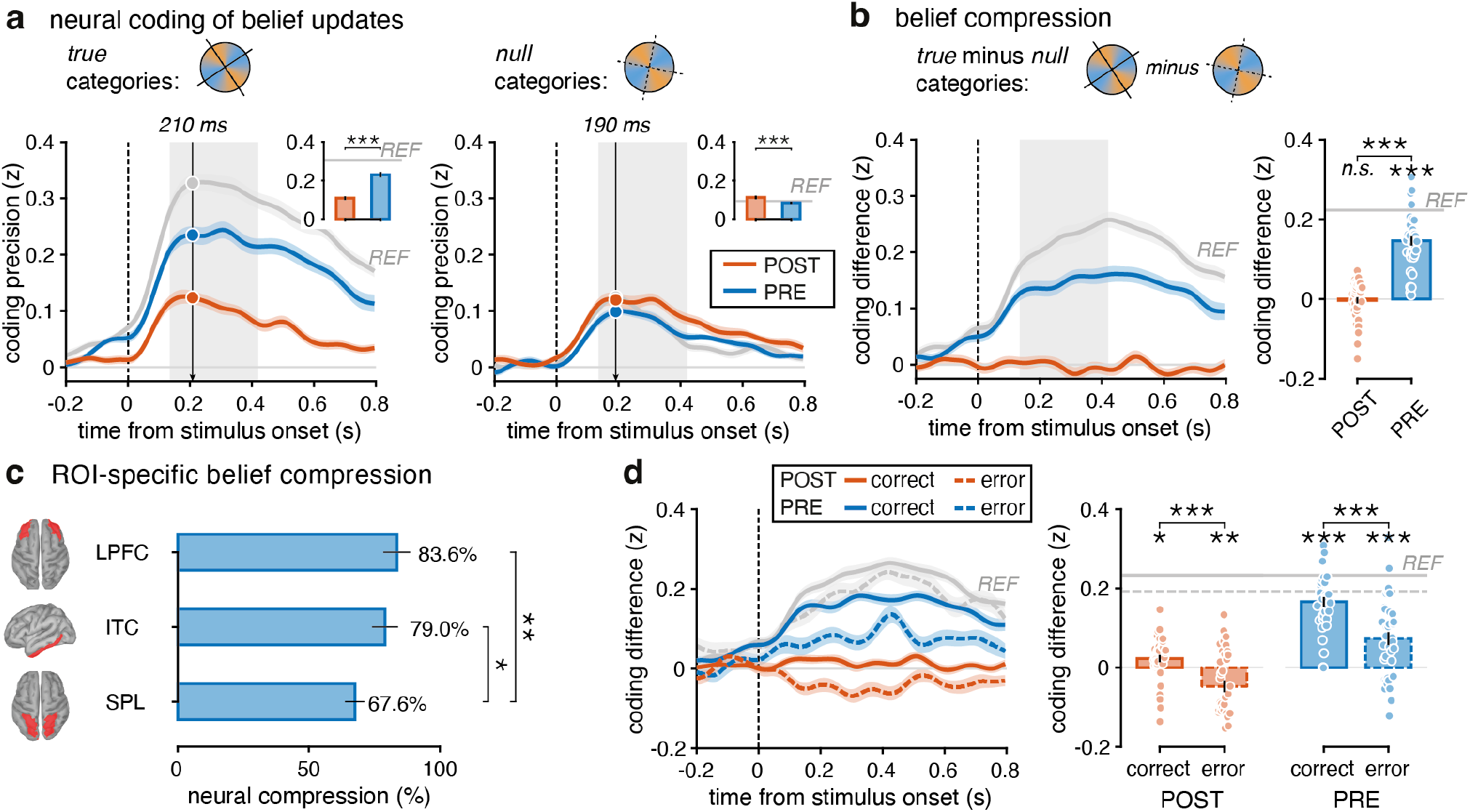
Compressed neural representations of decision-relevant information (N = 37). **(a)** Neural coding of belief update depending on category axis. Left: True belief update, computed by projecting the belief onto the true (decision-relevant) category axis, best decoded in REF and PRE but also decoded in POST. The coding precision reaches 90% of its peak value in at least one condition on [+135 +415] ms (gray shaded area; condition-specific: POST: [+135 +270] ms, PRE: [+150 +375] ms, REF: [+155 +415] ms). Right: Null belief update, computed by projecting the belief onto the null (decision-irrelevant) category axis along the category boundaries, better decoded in POST than in PRE. Top right insets: Coding precision averaged on the gray shaded area. **(b)** Compression on decision-relevant axis as difference between true and null belief update. Left: Time course of the compression. Right: Compression averaged on [+135 +415] ms. No difference from zero found in POST. In PRE and REF, the true belief update is much better represented than the null belief update. **(c)** Relative compression averaged on [+135 +415] ms for three brain regions in PRE: SPL, ITC and LPFC. The activity of each region reflects a high relative compression, even more for higher brain areas. **(d)** Neural coding of compression for correct (solid lines) and error (dashed lines) trials. Left: Time course. Right: Compression for correct and error trials averaged on the condition-specific time windows given in (a). The compression predicts participants’ accuracy in all conditions. In all panels: lines and shaded areas correspond to mean ± s.e.m., bars and error bars correspond to mean ± s.e.m. (jackknifed in (c)), small dots indicate single participants, three stars indicate a significant effect at *p* < 0.001, two stars at *p* < 0.01 and one star at *p* < 0.05, n.s. indicates no statistical difference.

We quantified neural compression as the difference in coding precision between true and null belief updates (**Fig. 5b** and **Supplementary Fig. 7a**). This analysis confirmed a significant compression in PRE (*t*_36_ = 12.8, *p* < 0.001), but not in POST, where evidence could only be accumulated in sensory space (*t*_36_ = −0.36, *p* = 0.71, BF_01_ = 5.30). We further compared compression in PRE across three brain regions in which belief updates could be decoded: the SPL, the inferotemporal cortex (ITC), and the lateral prefrontal cortex (LPFC; see **Methods** and **Supplementary Fig. 7b**). Relative compression – i.e., compression normalized by the coding precision of true belief updates – was stronger in the LPFC (83.6%) and ITC (79.0%) than in the SPL (67.6%) – a parietal region where evidence strength could also be decoded (jackknifed ITC-SPL *p* < 0.05 and LPFC-SPL *p* < 0.01; **Fig. 5c**).

Importantly, compression in PRE was stronger before correct decisions than before errors (one-tailed *t*_36_ = 5.44, *p* < 0.001; **Fig. 5d** and **Supplementary Fig. 8**). More surprisingly, fluctuations in neural compression around zero in POST also differed between correct decisions and errors (one-tailed *t*_36_ = 4.96, *p* < 0.001): although categories were unknown until the end of the trial, decision errors were preceded by a slightly *negative* compression, i.e., weaker coding of belief updates relative to the true than to the null categories. Together, these findings indicate that neural compression predicts accurate category-based decisions (in PRE), and that the inability to compress evidence on the decision-relevant axis is detrimental to decision accuracy (in POST).

### Stronger neural compression is associated with more precise inferences

By focusing limited resources on decision-relevant information, compression in category space might be an efficient strategy to compensate for the cost of imprecise inference. To test this hypothesis, we exploited individual differences and correlated each participant’s compression in PRE, measured from MEG activity (**Fig. 5**), with their inference imbalance and inference noise, estimated from behavior (**Fig. 3**). We found that neural compression did not correlate with inference imbalance (all uncorrected *p* > 0.05 after stimulus onset; **Fig. 6a**, left), but correlated negatively with inference noise (cluster-corrected *p* < 0.001; **Fig. 6a**, right). A multiple regression model including both parameters confirmed this selective relation between compression and inference noise (*β* = −0.696 ± 0.125, *p* < 0.001, estimate ± s.e.; unique contribution *r*^2^ = 0.476, *p* < 0.001; **Fig. 6b** and **Supplementary Fig. 9**).

**Figure 6.**
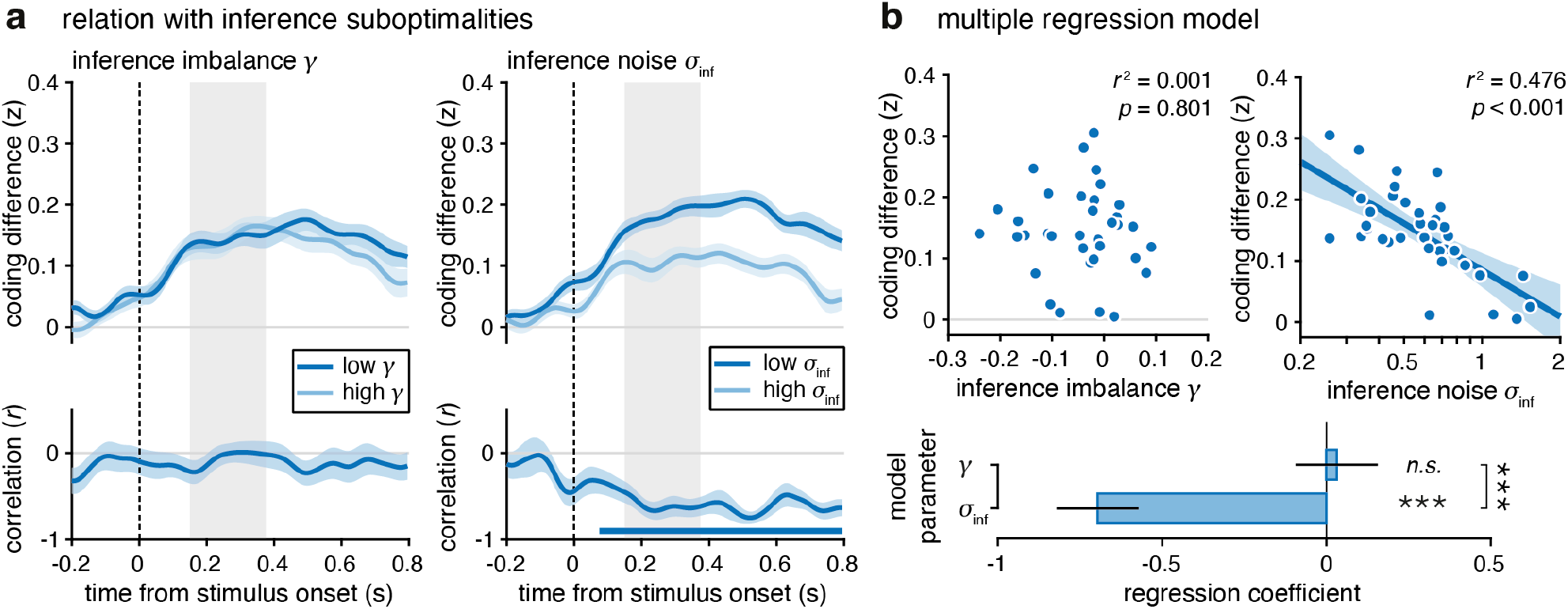
Inter-individual differences in neural correlates of compression in category space (N = 37). **(a)** Gradients in neural compression associated with inference suboptimalities in PRE. Top: Neural coding of compression when sorting participants in two median-split groups as a function of their best-fitting inference imbalance parameter *γ* (left) or inference noise parameter *σ*_inf_ (right). No qualitative relation found between *γ* and compression. Higher compression is associated with more precise inference (low *σ*_inf_, dark blue line), lower compression is associated with less precise inference (high *σ*_inf_, light blue line). The gray shaded area indicates the [+150 +375] ms time window, during which the coding precision of the true belief update reaches 90% of its peak value. Lines and shaded areas correspond to mean ± s.e.m. Bottom: Time-resolved correlation between participants’ inference imbalance parameter *γ* (left) and inference noise parameter *σ*_inf_ (right, log-transformed) and their compression. Higher compression is associated with smaller values of *σ*_inf_ from +95 ms after stimulus onset onward (cluster-corrected *p* < 0.001). Lines and shaded areas correspond to correlation and bootstrapped s.e.m. **(b)** Multiple regression model predicting individual neural compression (averaged over [+150 +375] ms) from participants’ inference imbalance parameter (left) and inference noise parameter (right). Dots indicate single participants, the line indicates significant partial regression controlling for the other parameter, shaded areas represent the 95% confidence interval. Bottom: Regression coefficients. Participants’ compression is predicted by their inference noise *σ*_inf_ but unrelated to their inference imbalance *γ*. Bars and error bars indicate estimate ± s.e., three stars indicate a significant effect at *p* < 0.001 and n.s. indicates no statistical difference.

We also assessed the relation between inference noise and the other stimulus characteristics decoded from MEG activity: stimulus orientation and evidence strength (**Supplementary Fig. 10**). Inference noise correlated negatively with the coding precision of both variables: weakly for stimulus orientation (*r*^2^ = 0.130, *p* = 0.029) and more strongly for evidence strength (*r*^2^ = 0.367, *p* < 0.001). A multiple regression model accounting for these correlations confirmed the selective relation between neural compression and inference noise (*β* = −0.515 ± 0.137, *p* < 0.001, estimate ± s.e.; unique contribution *r*^2^ = 0.186, *p* < 0.001). These findings indicate that stronger neural compression is associated with more precise inference.

### Neural compression makes RNNs more resilient to neural noise

We next sought causal evidence in artificial networks that stronger neural compression yields more precise inference. For this purpose, we trained 100 recurrent neural networks (RNNs) to perform the PRE and POST conditions of the task (see **Methods**). Each network had a single recurrent layer of 32 units. The category axis was provided to the networks as a noisy memory retrieval signal at each time step in PRE sequences, and as an exact post-sequence cue in POST sequences, mirroring the information available to human participants during accumulation (**Fig. 7a** and **Supplementary Fig. 11a**,**b**).

**Figure 7.**
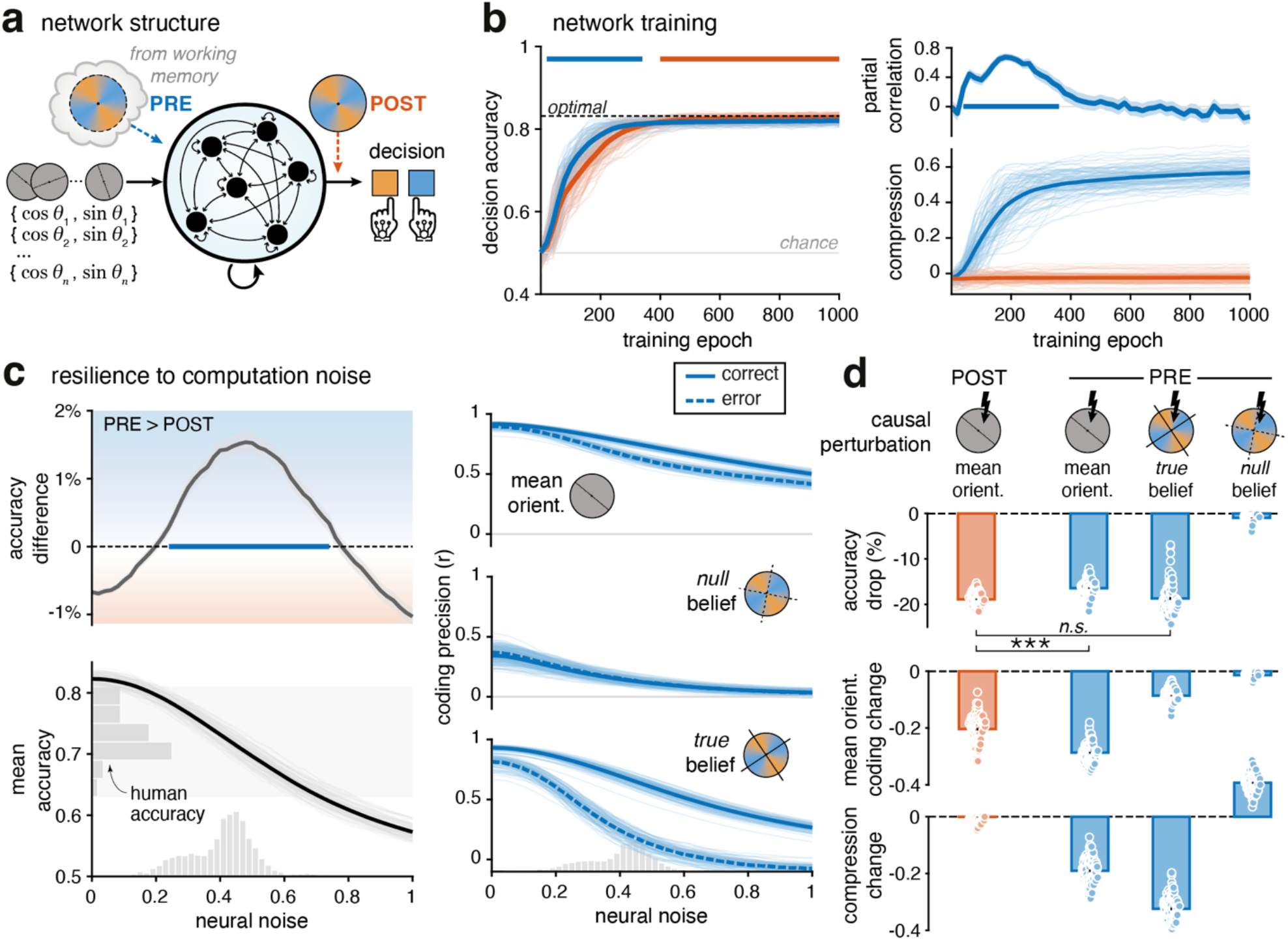
Benefits of compressing information in the presence of neural noise. **(a)** Network structure. At each time step, all 32 recurrent units receive the stimulus orientation (cos *θ*_*n*_, sin *θ*_*n*_). In PRE trials, the units receive the noisy memory retrieval of the category axis together with the stimulus orientation. The exact category axis is provided at the end of POST sequences. **(b)** Network training (N = 100). Left: Networks trained simultaneously on PRE and POST trials perform the categorization task near-optimally after 1,000 epochs (dashed line: Bayes-optimal accuracy), learning PRE faster (epochs 20-340) but reaching a slightly higher final accuracy in POST (epochs 400-1,000; both cluster-corrected *p* < 0.001). Right: Compression decoded from recurrent units during PRE and POST trials (bottom) and its correlation with decision accuracy along training when controlling for the decodability of the sequences’ mean orientation (top). Compression and accuracy are highly correlated during the steep learning phase (epochs 40-360, cluster-corrected *p* = 0.002). **(c)** Resilience to neural noise. Left: Effect of noise added to recurrent computations on mean accuracy across PRE and POST (bottom) and on the PRE-POST accuracy difference (top). Human PRE and POST mean accuracy is indicated by the left gray histogram. PRE sequences are better categorized than POST sequences for noise levels 0.22-0.78 (one-tailed cluster-corrected *p* < 0.001; bottom gray histogram: human-matched noise levels). Right: Effect of noise on the coding precision of the mean orientation (top), null belief (middle) and true belief (bottom) for correct (solid lines) and error (dashed lines) trials from PRE, with the same bottom gray histogram. True belief coding precision strongly predicts decision accuracy under noise. **(d)** Causal benefit of neural compression under noise. Effect on accuracy (top) of selectively perturbing the coding of the mean orientation in POST, and of the mean orientation, true belief and null belief in PRE. Middle and bottom: Change in mean orientation coding and in compression. Perturbing true belief, but not the mean orientation, eliminates the PRE advantage over POST. In all panels: thick lines indicate means across networks, thin lines and dots correspond to individual RNNs, bars and error bars correspond to mean ± s.e.m., three stars indicate a significant effect at *p* < 0.001 and n.s. indicates no statistical difference.

The networks were trained simultaneously on both PRE and POST sequences and reached near Bayes-optimal accuracy after 1,000 epochs (**Fig. 7b**, left), with PRE sequences learned faster than POST sequences (cluster-corrected *p* < 0.001, Cohen’s *d* = 0.89) and a slightly higher final accuracy in POST than in PRE (POST: 82.86%; PRE: 82.02%; cluster-corrected *p* < 0.001, Cohen’s *d* = 1.33). We then decoded three variables from recurrent-layer activity: 1. the mean orientation of the sequence, 2. the accumulated belief relative to the true categories, and 3. relative to the null categories. As for MEG activity, we quantified neural compression as the difference in coding precision between true and null beliefs. Unlike for MEG activity, we decoded accumulated beliefs rather than belief updates, since RNNs give direct access to the activation of every recurrent unit. RNNs compressed beliefs in PRE but not in POST trials, and the strength of compression mirrored the steep rise in accuracy early in training, even when controlling for the decodability of the mean orientation (mean partial *r* = 0.47, cluster-corrected *p* = 0.002; **Fig. 7b**, right). Compression did not imply that sensory information was discarded: PRE networks still coded the mean orientation of the sequence (**Fig. 7c**).

Because human evidence accumulation was corrupted by substantial inference noise, we probed the resilience of these RNNs to isotropic (non-directional) noise injected into their recurrent layer (see **Methods**). Neural noise reduced mean accuracy in both conditions, yet networks performed more accurately on PRE trials than on POST trials at noise levels matching human performance (noise-matched range, 5th-95th percentile: [0.21 0.56], significant cluster: [0.22 0.78], Cohen’s *d* = 0.99, one-tailed cluster-corrected *p* < 0.001; **Fig. 7c**, left). Although this accuracy advantage was smaller than in humans (1.25%, compared to 4.3% in humans), it suggests that the compression strategy that emerges in PRE confers greater noise resilience than accumulating the mean orientation of the sequence in sensory space, as in POST. To identify which decoded variable best accounts for this advantage, we asked which one most strongly predicted correct decisions under increasing noise (**Fig. 7c**, right). Errors were associated with slightly worse decoding of the mean orientation (mean correct-error difference: Δ*r* = 0.08) but with a larger drop in decoding of the true category belief across noise levels (mean correct-error difference: Δ*r* = 0.36), consistent with the idea that compressed category-space representations, rather than sensory-space accumulation, support accurate decisions under noise.

To test whether this advantage is specific to recurrent noise, we also injected noise into the network input (see **Methods**). Input noise reduced accuracy in both conditions but produced no PRE accuracy advantage over POST (**Supplementary Fig. 12**), indicating that compression specifically protects against noise arising within recurrent computations, not against noise arising from upstream computations (or from external noise in the stimulus itself).

To establish a causal link between compression and noise resilience, we selectively disrupted the coding of each strategy-relevant variable in the recurrent layer (**Fig. 7d** and **Supplementary Fig. 13**; see **Methods**). Because accumulating the mean orientation in sensory space is the only strategy available in POST, perturbing its coding in POST sets a reference accuracy drop (−18.9%). We compared this reference with the accuracy drops produced in PRE trials by perturbing the coding of mean orientation, true belief, or null belief. Perturbing mean orientation coding in PRE produced a significantly smaller accuracy drop than in POST (PRE: −16.4%; *t*_99_ = −14.31, *p* < 0.001). By contrast, perturbing true-belief coding in PRE produced an accuracy drop (−18.7%) similar to the POST reference (*t*_99_ = −0.54, *p* = 0.58, BF_01_ = 7.82), indicating that PRE decisions rely more on the compressed belief representation than on the coding of mean orientation (**Fig. 7d**, top). Perturbing mean orientation coding in PRE also reduced neural compression (**Fig. 7d**, bottom), suggesting that part of its effect on accuracy was mediated by the compressed belief representation. Perturbing null-belief coding had a negligible effect on accuracy, ruling out that any disruption of category-space representations would produce this effect. Perturbing true-belief coding strongly reduced compression but only weakly affected the coding of mean orientation, whereas perturbing null-belief coding increased compression without affecting the coding of mean orientation (**Fig. 7d**, middle and bottom; **Supplementary Fig. 13**), confirming the specificity of these two perturbations. Together, these results identify compressed category-belief representations as the key mechanism underlying noise-resilient inference in PRE trials in RNNs.

## Discussion

Evidence accumulation is widely seen as the core process for making accurate perceptual decisions under uncertainty. Surprisingly, the representation space in which humans accumulate evidence during simple categorical decisions has long remained unclear. Here we used a visual categorization task that manipulates whether participants can accumulate evidence in sensory space or in category space. We found that human observers preferentially accumulate compressed evidence in an abstract category space, by projecting momentary sensory information onto the decision axis, even though this axis changed from one trial to the next. When forced to accumulate evidence in sensory space, the same participants did so with more inference noise, a stronger inference imbalance (i.e., more leaky accumulation), and delayed responses in the presence of decision-irrelevant evidence. MEG recordings revealed neural signatures of evidence compression whose strength predicted decision accuracy. RNNs trained on the same task were more resilient to noise in their recurrent computations when using this compression strategy.

An important feature of our task is that neither representation space confers a theoretical advantage in terms of ideal accuracy^3^: projecting each stimulus onto the decision axis before accumulation (category space) or projecting the accumulated sensory evidence onto the decision axis after accumulation (sensory space) both predict identical behavior. Given this formal equivalence, why do human observers preferentially accumulate evidence in category space? A first reason could be that accumulating evidence in category space requires updating only a single scalar log-posterior odds ratio, whereas accumulating evidence in sensory space requires updating a posterior distribution over orientation (at the very least, a two-dimensional vector of sufficient statistics). If the computational resources available for evidence accumulation are limited – e.g., through bounded firing rates or a limited number of neurons – this difference in dimensionality can affect the precision of inference. Efficient coding principles, initially proposed for sensory systems^25^ and more recently extended to decision variables such as subjective value^34^, posit that neural systems allocate limited coding resources so as to maximize the information they convey. In our task, the information that matters for the decision is the projection of each stimulus onto the decision axis. Encoding a single, decision-relevant variable allows the full dynamic range of the neural population to be devoted to it, thereby increasing its signal-to-noise ratio compared with a representation that must share the same resources across two dimensions. Consistent with this account, participants showed less inference noise and a weaker inference imbalance when they could accumulate evidence in category space (in PRE and REF). Moreover, RNNs with bounded units were more resilient to noise arising in their recurrent computations, but not in their inputs, when they compressed evidence. A second (non-exclusive) reason is that accumulating evidence in category space allows projecting away decision-irrelevant information – i.e., the component of each stimulus that is orthogonal to the decision axis, together with internal noise along this dimension – that may otherwise interfere with the processing of decision-relevant information. In line with this hypothesis, the decision-irrelevant evidence provided by the sequence slowed down responses in POST, but not in PRE and REF (**Supplementary Fig. 2d**). Nonlinear attractor models^13,35^ show how recurrent dynamics can make relevant and irrelevant inputs interact during evidence accumulation, producing interference patterns that are not predicted by linear, optimal accumulation models.

Our findings relate to previous work showing that encoding strategies that deviate from optimal evidence accumulation can nevertheless increase decision accuracy when decisions are corrupted by internal noise. In particular, discounting weaker or outlying samples during integration – termed ‘selective integration’^36^ and ‘robust averaging’^37^ – and overweighting larger magnitudes during numerical comparison^38^ have been shown to protect decisions from inference noise. Compressing evidence onto the decision axis can be seen as such a noise-resilient cognitive strategy, by restricting evidence accumulation to the decision-relevant dimension. But unlike these strategies, compression does not deviate from ideal evidence accumulation, as accumulating evidence in category space is sufficient to perform Bayes-optimal categorization.

The evidence strength provided by each stimulus could be decoded from MEG activity in PRE even though the stimulus-category mapping changed from one trial to the next (**Supplementary Fig. 1**). Because the same decoder was applied across trials with different mappings, this neural code is abstract with respect to the orientations that define the categories on each trial: it reflects how strongly each stimulus favors either category, irrespective of its orientation. This abstract format echoes the coding of learned visual categories by neurons in the monkey prefrontal^39^ and lateral intraparietal^23^ cortices, whose activity reflects category membership rather than the visual features of the stimuli. It is also compatible with empirical evidence for an abstract neural code for evidence accumulation across different sensory features^21,40–44^. Unlike in these studies, however, the category boundary changed on every trial in our task, which required participants to flexibly re-map sensory evidence onto a new decision axis on each trial.

Compared with POST, accumulating evidence in category space in PRE simultaneously enhanced the coding of belief updates relative to the true categories and reduced the coding of belief updates relative to the null categories. This dual tuning through amplification of decision-relevant information and reduction of decision-irrelevant information is constitutive of a ‘compressed’ neural representation of evidence. Importantly, this compression did not come at the expense of early sensory processing, since the early coding of stimulus orientation was unaffected in PRE. This pattern is reminiscent of two previous findings. First, the monkey prefrontal cortex represents both relevant and irrelevant sensory inputs during context-dependent decisions, with relevant inputs selected through recurrent dynamics during evidence accumulation^42^. Second, task-irrelevant stimulus dimensions are compressed in humans and in neural networks trained on context-dependent tasks^45^. That this compression covaried with decision accuracy – both within and across participants – is consistent with its relevance for accurate decisions. Although participants appeared to accumulate evidence in category space when categories were provided before the sequence in PRE, the coding of evidence strength and belief updates appeared less precise in PRE than in REF (**Figs. 4b** and **5a**). This difference likely arises because participants had to maintain the category orientations in working memory in PRE, whereas these were continuously displayed in REF. Indeed, working memory representations of fine-grained features such as orientations have limited precision during sustained maintenance^46^. Behaviorally, the accuracy difference between PRE and REF can be accounted for by an imprecision of about 15° (s.d.) in the maintenance of the category axis (**Supplementary Fig. 11a**).

Our behavioral and neural findings in humans are correlational: they show that compression is associated with more precise inference, but not that it causes it. We therefore trained RNNs on the same task to test whether compression makes decisions more resilient to internal noise. RNNs spontaneously compressed evidence along the decision axis in PRE and were more robust to noise in their recurrent computations when doing so, providing causal evidence (albeit in artificial networks) that this strategy improves decision-making under uncertainty. Importantly, computation noise in RNNs does not merely act as a constraint but actively shapes their computational dynamics, enabling adaptive behaviors that generalize beyond training conditions^47^. A growing body of work uses RNNs to model cognitive processes descriptively^48,49^ by fitting individual choices and then investigating subject-specific strategies. Here, instead, we used RNNs not to fit behavior but to probe the computational advantages of a particular strategy. Recent studies have also started to use RNNs this way to better understand the nature of animal and human abilities^47^ and idiosyncrasies^50^.

Together, these findings reveal a representational strategy in which sensory evidence is flexibly accumulated according to its relevance for the upcoming categorical decision. In humans, this neural compression strategy is associated with reduced inference noise; in RNNs, it increases resilience to such noise. Our multimodal combination of behavioral, MEG, and neural-network analyses shows that this neural compression is both reflected in human brain activity and beneficial for performance in artificial neural networks subject to internal noise. Examining not just how evidence is processed but why humans favor specific cognitive strategies provides insight into the principles that guide adaptive decision-making.

## Methods

### Participants

Thirty-seven healthy adult participants between 20 and 35 years old (18 females, age: 24y5m ± 3y11m) with normal or corrected-to-normal vision took part in the study. The recruitment took place between March and August 2021 and participants were screened for the absence of any history of neurological and psychiatric disease or any current psychiatric medication. All participants provided written informed consent and received 90 euros as compensation for their participation. The study followed guidelines from the Declaration of Helsinki and specific procedures which received ethical approval from relevant authorities (Comité de Protection des Personnes île-de-France VI, ID RCB: 2017-A01778-45).

### Experimental task

The task was a probabilistic cue combination task corresponding to a modified version of the weather prediction task^7,51^ framed as a card game (**Fig. 1b**). On each trial, a sequence of 4, 8 or 12 cards was drawn from one of two decks (blue or orange category) and the participant had to guess from which deck the sequence was drawn. Each card depicted a circle containing an oriented black bar sampled from a circular Gaussian distribution (von Mises distribution). Both von Mises distributions underlying the blue and orange categories had the same fixed concentration parameter *κ* = 0.5 but different means, separated by an angle of *π*⁄2. The means varied pseudo-randomly along the circle across trials. Participants had to accumulate the evidence provided by each card to infer the most likely category of the sequence. A category-mapping cue, consisting of a colored disc displaying an angular gradient between orange and blue (through gray) – the two cardinal colors being spaced by *π*⁄2 – allowed participants to relate the orientation of the cards to the two categories.

To identify the representation used to accumulate evidence, we manipulated when the category-mapping cue was presented across three conditions. In the post-mapping condition (POST), the cue was presented after the sequence, so that participants had no knowledge of the categories during evidence accumulation. In the pre-mapping condition (PRE), the cue was presented before the sequence, so that participants could use their knowledge of the categories during evidence accumulation. In the reference condition (REF), the cue was displayed throughout the trial: before, during and after the sequence.

Each trial started with a cue card presented for exactly 0.5 s, followed by an empty card for ~0.5 s; the sequence was then presented at an average rate of 3 Hz with each card being shown for exactly 100 ms. After the sequence, an empty card appeared for ~0.5 s followed by a response probe card indicating the response mapping (left or right button for the orange or blue category), which served as the ‘go’ signal for the response. In the POST condition, the cue card was an orange-blue scrambled noisy Gabor patch matching the energy and luminance of the category-mapping cue. In the PRE and REF conditions, the cue card was the category-mapping cue. In PRE, the ‘go’ signal card indicated only the response mapping but did not show the category-mapping cue. In POST and REF, the response probe card showed both the response mapping and category-mapping cue.

The length of the sequence was sampled pseudo-randomly and uniformly across trials, such that participants could not anticipate when they were prompted to respond. The response mapping was randomized across trials. Participants responded by pressing one of two buttons on a Cedrus response pad (Cedrus Corporation) with their left or right index finger, using the indicated response mapping. They had no limitation in time to respond. After each trial, they received visual feedback in the form of a ‘+’ sign for positive feedback or a ‘×’ sign for negative feedback, appearing briefly (250 ms) in the center of the fixation point.

The task was divided into six blocks of 90 trials, each lasting ~10 min. The three conditions were interleaved across trials and fully matched per participant, resulting in 180 trials per condition per participant. The first half of the experiment was repeated in the second half so that, for the same condition, each sequence was presented twice. Participants were familiarized with the task during a training session without MEG recordings, on average 2 weeks earlier, responding on a regular keyboard. During training, they were instructed explicitly that the decks were partially shuffled, such that the orange deck contained mostly orange cards but also blue cards, and the blue deck contained mostly blue cards but also orange cards. They ran 2 blocks of 90 trials each, the first block difficulty was reduced by increasing the von Mises concentration parameter *κ* to 1. The second block had the same difficulty and same length as an experimental block (*κ* = 0.5, *n* = 90 trials). Performance in these practice blocks was used to verify participants’ understanding of the task before the MEG session. All trained participants qualified to proceed to the MEG experiment. Before the MEG session, participants had to re-read the written instructions, but no additional training block was performed.

All stimuli were presented on a gray background, at a viewing distance of 80 cm. Visual fixation was monitored online throughout the main task using an EyeLink 1000 eye-tracking system (SR Research), using monocular tracking of the dominant eye at a sampling frequency of 1,000 Hz. MEG data were recorded throughout the experiment.

### Behavioral analyses

We computed the Bayes-optimal accuracy as the fraction of sequences whose log-likelihood ratio was in favor of their generative category. To estimate the sensitivity of participants’ decisions to the evidence provided by each sequence, we regressed their choices against the log-likelihood ratio of the sequence using a logistic regression, and used the slope of this regression as the sensitivity coefficient (**Fig. 2b**). We used a similar approach to estimate decision suboptimality for each sequence length: the inverse of the slope of a probit regression of choices against the log-likelihood ratio of the sequence corresponds to the standard deviation of the deviations from Bayes-optimal decisions (**Fig. 2c**). The resulting variance was compared to the prediction of a model with inference noise only, in which this variance increases linearly with the number of stimuli. Finally, we estimated the integration kernel by regressing participants’ choices against the log-likelihood ratio of each of the first four and last four stimuli of the sequence (probit regression, the other stimuli being summed in a single regressor; **Fig. 2d**).

### Bias-variance decomposition

We followed the method described in previous work^7^ based on choice consistency. In practice, we used the best-fitting inference noise parameter found to match participants’ choice variability (**Fig. 2c**) to simulate responses. We allowed the noise to be correlated and progressively increased this level of noise correlation, therefore increasing systematicity in predictions. We computed the fraction of consistent choices for each simulation, and could then estimate the level of correlation matching each participant’s fraction of consistent choices. Correlated ‘noise’ corresponds to the bias term, while uncorrelated noise corresponds to the variance term: genuine inherent stochasticity in the computations.

### Computational model

We implemented a suboptimal noisy inference model based on the normative accumulation model for two alternatives^2^, assuming equal prior probabilities for the two categories. For a given sequence of orientations *θ*_1_, …, *θ*_*n*_, a Bayes-optimal observer accumulates the evidence provided by each stimulus expressed as log-likelihood ratio ℓ_*i*_ = log(*p*(*θ*_*i*_|*A*)⁄*p*(*θ*_*i*_|*B*)) to form the log-posterior belief 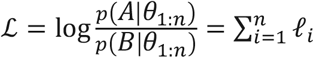 and chooses the category (*A* or *B*) based on the sign of ℒ (*A* if ℒ > 0, *B* otherwise). The likelihood of sampling *θ*_*i*_ from category *A* with generative mean *µ*_*A*_ is given by the von Mises distribution with fixed concentration *κ*, adjusted for the half-circle symmetry of the task where each orientation is defined modulo *π*, so that:

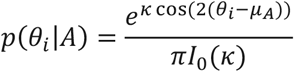

where *I*_0_(·) is the modified Bessel function of order 0. Given that the categories *A* and *B* are orthogonal *µ*_*B*_ = *µ*_*A*_ + *π*⁄2, the log-likelihood ratio for the *i*-th stimulus simplifies to:

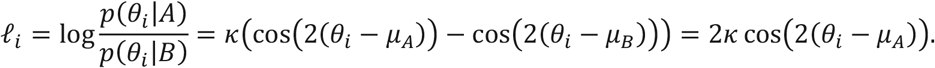

We introduced suboptimality in the normative accumulation process through three optional parameters and a lapse rate:

1. a time-invariant exponential inference imbalance *γ* capturing the unequal weighting of the previous log-posterior relative to the new evidence:

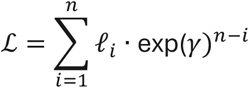

for *γ* < 0, past observations are down-weighted, similarly to an inference leak, since exp(*γ*) < 1, producing a recency effect; *γ* > 0 leads to a primacy effect by putting more weight on early observations and *γ* = 0 corresponds to a Bayes-optimal weighting of prior and likelihood.
2. inference noise *σ*_inf_ capturing variability at the inference stage through noisy log-likelihood ratios:

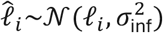
3. selection noise *σ*_sel_ capturing variability at the selection stage through noisy log-posteriors:

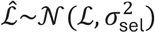

and the chosen category (*A* or *B*) is based on the sign of 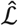. This parameter was ruled out by model selection.

The lapse rate *ε* at the selection stage captured occasional attentional lapses or errors during choice, irrespective of the posterior belief.

This category-space model accumulates already categorized evidence, but it is mathematically equivalent to a sensory-space model that projects the accumulated resultant vector onto the category axis at the end of the sequence (see Supplementary Information). We therefore used this model in all conditions.

### Fitting procedure

We implemented this canonical model with distinct inference imbalance, inference noise and selection noise parameters for each condition and a common lapse rate parameter across the three conditions, resulting in ten free parameters. Each parameter except the lapse rate could be enabled (free) or disabled (KO), resulting in several model variants. Each variant was fitted to participants’ choices. Per-trial choice log-likelihoods were estimated using a particle filter and then combined with prior distributions over parameters to obtain the log-posterior. The prior distributions over parameters were the same across conditions and were defined as truncated normal, exponential and beta distributions (inference imbalance *γ*: normal distribution with mean 0 and s.d. 0.2, range [−2 2]; inference noise *σ*_inf_: exponential distribution with mean parameter *µ* = 0.5, range [0 5]; selection noise *σ*_sel_: exponential distribution with mean parameter *µ* = 1, range [0 5]; lapse rate *ε*: beta distribution with shape parameters *α* = 1 and *β* = 49, range [0 1]). The unnormalized log-posterior was passed to the Variational Bayesian Monte Carlo (VBMC) algorithm^52,53^ (version 1.0.12) returning a variational approximation of the full posterior and a lower bound on the log-marginal likelihood. We took the posterior mean (and standard deviation) to obtain best-fitting parameter values (and standard deviation). We used the lower bound on the log-marginal likelihood as model evidence for Bayesian Model Selection (BMS) analysis. BMS used the random-effects Dirichlet approach implemented in SPM12 with the default Dirichlet prior set to one to estimate the distribution over model variants. We then grouped model variants according to the presence or absence of a given suboptimality parameter (e.g., inference imbalance / inference noise / selection noise: “free” or “KO”; lapse rate always “free”) and ran factorial family-wise model comparisons per condition (**Fig. 3b**). We estimated family exceedance probabilities based on Dirichlet sampling (10^6^ draws). Finally, we ran model simulations based on best-fitting model parameters to qualitatively validate the goodness-of-fit of the selected model.

### Model and parameter recovery

Before fitting human choices, we verified that model comparison could arbitrate between the presence or absence of the possible sources of suboptimality by generating synthetic data from each parameter combination and verified that they recovered the correct source of suboptimality. We ran model recovery considering variants for one condition – and letting the parameters for the other conditions always free – model variants had therefore only four parameters (*γ, σ*_inf_, *σ*_sel_: free or KO and *ε* always free resulting in eight model variants). The parameters used to generate the synthetic data were sampled from their prior distributions as previously defined. For each model variant, we generated 37 synthetic datasets using the stimulus sequences seen by each of the 37 participants and the resulting simulated choices were fitted by all possible model variants. We used the Dirichlet parameterization as described above to estimate the distribution over recovered model variants. We then sampled 10^6^ times from the estimated Dirichlet distributions and computed the family-wise exceedance probability for each source of suboptimality. All sources of suboptimality were successfully recovered with exceedance *p* > 0.999 (**Supplementary Fig. 4a**, left). We also verified the ability of the model to provide accurate estimates of each parameter through parameter recovery by correlating each best-fitting parameter to the parameter values used for simulation. All recovered parameters were highly correlated to simulation parameters except for the lapse rate parameter (inference imbalance *γ*: *r*^2^ = 0.932; inference noise *σ*_inf_: *r*^2^ = 0.830; selection noise *σ*_sel_: *r*^2^ = 0.711; lapse rate *ε*: *r*^2^ = 0.272, all *p* < 0.01; **Supplementary Fig. 4a**, right).

### Magnetoencephalography

MEG data were recorded using a whole-head Elekta Neuromag TRIUX system (Elekta Instrument AB, Stockholm, Sweden) composed of 204 planar gradiometers and 102 magnetometers, at a sampling frequency of 1,000 Hz. Prior to the experiment, we digitized each participant’s head shape in the MEG coordinate frame and used four additional head position indicator (HPI) coils – whose positions were also digitized – to monitor and correct for small head movements across blocks.

We preprocessed MEG signals as follows: we removed manually detected non-physiological jumps, then used temporal Signal Space Separation (tSSS) to remove magnetic noise from external sources. Then, we used a supervised Principal Component Analysis (PCA) procedure to correct for stereotyped ocular and cardiac artifacts. The onset of artifacts (eye blinks or cardiac R-peaks) was automatically detected from the auxiliary electrodes (EOG and ECG) and then synchronized with MEG signals using a threshold-based approach. The MEG signals were epoched from −100 ms to +400 ms around artifact onset, and PCA was applied to these epochs to extract the spatial components of ocular and cardiac artifacts. We removed the first component of the ocular artifact and the first and second components of the cardiac artifact from the continuous MEG signal. Finally, MEG signals were high-pass filtered at 0.5 Hz, low-pass filtered at 40 Hz, down-sampled to 200 Hz and epoched from −200 to +800 ms around stimulus onset. We applied a linear detrending and a low-pass filter to the epoched MEG signals, whose frequency depended on the analysis (16 Hz for stimulus orientation decoding, 8 Hz for the other variables). We then computed their analytic representations using the Hilbert transform, resulting in twice the number of MEG signals (408 from planar gradiometers and 204 from magnetometers). This preprocessing pipeline was implemented using the FieldTrip toolbox^54^ and additional custom scripts written in MATLAB.

### Multivariate pattern analyses of MEG signals

We applied multivariate pattern analyses (MVPA) to each time sample of the stimulus-locked MEG epochs (1440 per participant and per condition) to estimate the neural patterns associated with different characteristics of each stimulus *i* within a given sequence. The first one was its orientation *θ*_*i*_ described by cos(2*θ*_*i*_) and sin(2*θ*_*i*_); the second was its evidence strength, described by the magnitude of its log-likelihood ratio |ℓ_*i*_| = |2*κ* cos(2( *θ*_*i*_ – *µ*_*A*_))|; the third was the belief update triggered by the stimulus, defined as its log-likelihood ratio ℓ_*i*_ signed by the belief accumulated so far sign(ℒ_*i−1*_). Positive values correspond to belief-consistent evidence whereas negative values correspond to belief-conflicting evidence. The belief accumulated before each stimulus was estimated with a particle filter simulating the best-fitting model of each participant, conditioned on the participant’s choice. These category-based variables are computed with respect to the true generative orientation *µ*_true_ = *µ*_*A*_ or *µ*_*B*_ (and can be simplified to only being expressed by *µ*_*A*_ as seen above), but can also be computed with respect to any axis, especially when the category-mapping cue is unknown during accumulation as in POST. As a baseline for assessing compression, we used the least decision-relevant axis, i.e., the null categories corresponding to the boundary between the orange and blue categories *µ*_null_ = *µ*_*A*_ + *π*⁄4. The null log-likelihood ratio for the *i*-th stimulus becomes 2*κ* cos(2( *θ*_*i*_ − *µ*_null_)) and the null belief update is this null log-likelihood ratio signed by the already accumulated belief with respect to the null category. These stimulus characteristics showed minimal correlation with each other for a given stimulus *i* and between successive stimuli *i* and *i*+1 inside a sequence (all *r*^2^<0.05). The decoding pipeline consisted of predicting each of these variables from the Hilbert-transformed MEG neural signals of the gradiometers at each time step using ridge regression (with regularization factor set to 500) with 10-fold cross-validation across epochs. The decoding precision was computed as the Fisher transform of the Pearson correlation coefficient between the true and predicted values on the test epochs, resulting in a normally distributed variable. For stimulus orientation, the decoding precision was averaged across the cosine and sine decoders. Decoding precision was also computed separately for correct and error trials, defined as choices consistent or not with the sign of the log-likelihood ratio of the sequence (i.e., with the Bayes-optimal choice), from the same cross-validated predictions.

### MEG source reconstruction

Participants’ anatomical magnetic resonance images were acquired with a Verio 3T scanner (Siemens AG, Munich, Germany) and segmented with the FreeSurfer ‘recon-all’ pipeline^55^. We used Brainstorm^56^ and followed their tutorial pipeline for subsequent analyses: anatomical surfaces were down-sampled to 5003 vertices and re-aligned to MEG spatial coordinates using participants’ digitized head shape and fiducial markers. The cortex surface was modeled with overlapping spheres. Noise covariance matrices were computed on a separate empty-room recording lasting approximately 3 minutes acquired after each participant’s session. We estimated the activity at the source level using the minimum-norm imaging method, and the current dipole models were assumed to be oriented normally to the cortex surface. Each resulting individual source map was projected on a template anatomy (ICBM152), allowing for comparison and averaging across participants. The searchlight-based MVPA at the source level was conducted before projecting on the default anatomy with a sphere containing each vertex itself and its 9 closest neighbors in Euclidean distance. We conducted neural compression analyses on three pre-defined regions of interest (SPL, ITC, LPFC, averaged over both hemispheres) using parcellations from the Desikan-Killiany atlas after projection on default anatomy (for LPFC, we used the rostral middle frontal parcellation from the atlas). At the ROI level, we computed the relative neural compression as the difference between true and null belief update decoding divided by the true belief update and computed the jackknife mean.

### Recurrent Neural Networks

We implemented standard (Elman) Recurrent Neural Networks consisting of a single recurrent layer of 32 units to perform the categorization task. Each time an orientation *θ* is presented in a sequence *s*, the network updates its hidden state by combining the current input *X*_1_ with the previous hidden activity *Z*_*θ* −_ through linear transformations (weight matrices ***W***_*ih*_ and ***W***_*hh*_ plus bias ***B*)**, followed by a hyperbolic tangent nonlinearity, and the network output is computed with a linear readout (output weight matrix ***W***_*fo*_) at the end of the input sequence:

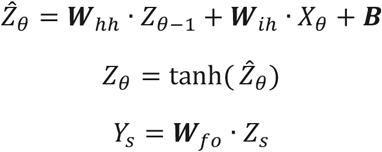

Each input *X*_*θ*_ was five-dimensional, containing the cos and sin of a noisy memory retrieval of the true category axis in PRE (set to zero for POST), the cos and sin of each presented orientation and a ‘go’ signal set to 1 only at the last time step and 0 otherwise. Each input sequence of *n* orientations had length (*n*+1), one step per orientation plus a final step containing the post-sequence category information, i.e., the exact category axis in POST (set to zero for PRE), and the ‘go’ signal set to 1. In PRE sequences, the noisy memory retrieval of the true category axis was sampled at each time step from a Gaussian distribution centered on the true axis with a standard deviation of 15° matching participants’ accuracy in PRE (**Supplementary Fig. 11a**).

Input-to-hidden weights were initialized using Xavier normal initialization^57^, with gain adapted for the tanh activation function, and hidden-to-output weights with the default gain. Recurrent weights were initialized orthogonally^58^. Recurrent biases were initialized to zero. Hidden states were initialized to zero at the start of each sequence. All networks were trained with the Adam optimizer on a binary cross-entropy loss with logits, computed from the output after processing of the full input sequence. Training was performed in batches of 512 orientation sequences per sequence length from 2 to 12 (unlike humans performing the task only on sequences of length 4, 8 or 12), so that the optimizer was updated after having seen 512 × 11 = 5,632 different sequences. The sequence category axes were generated uniformly. We trained 100 networks in both PRE and POST conditions (each corresponding to half of the batch) using a learning rate of 0.001 and stopping after 1,000 epochs (one update per epoch, on newly generated sequences).

Neural noise was introduced by adding Gaussian noise with standard deviation *σ* to the pre-activation of each recurrent unit at each time step,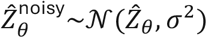. We additionally tested the effect of input noise by adding normally distributed noise to each sequence orientation *θ* with standard deviation from 0 to 40°. To compare networks and humans, we computed, for each participant and each network, the recurrent noise level at which the accuracy of the network (averaged across PRE and POST) matched that of the participant; the noise-matched range reported in the Results corresponds to the 5th-95th percentiles of these noise levels.

Similar to MEG analyses, we applied multivariate pattern analyses to the activity of trained units of the recurrent layer. We decoded cumulative sequence characteristics in these artificial agents: the mean orientation and the belief (accumulated evidence computed with respect to the true category axis or to the null category axis), and computed belief compression as the difference between the true and null beliefs. Trained networks were tested on 1,000 new sequences of 4, 8 and 12 stimuli in each condition, and each variable was decoded from the hidden activity after each stimulus using ridge regression (as for MEG), with decoding precision measured as the Pearson correlation coefficient (for the mean orientation, averaged across the decoders of the cumulative cosine and sine of the orientations). Decoding precision was also computed separately for correct and error sequences (relative to the generative category), from the same cross-validated predictions.

To causally disrupt the neural coding of these variables, we identified the direction in recurrent activation space most predictive of each variable using ridge regression decoding weights. At each time step, we injected normally distributed noise into the pre-activations of recurrent units along the identified direction. Importantly, to ensure comparable perturbation strength across variables, the noise amplitude was normalized by the standard deviation of the pre-activations projected onto each direction. The mean orientation perturbation was defined as the average of independent perturbations applied along the cosine and sine orientation directions. All perturbation conditions were applied to identical sequences, differing only in the direction along which selective noise was injected. Perturbation strengths, i.e., the noise standard deviation relative to the standard deviation of the projected pre-activations, ranged from 0 to 5, and their effects were averaged across strengths; accuracy drops are expressed as a percentage of the unperturbed accuracy.

### Statistical testing and reproducibility

Unless noted otherwise, statistical analyses of differences between scalar metrics relied on two-tailed parametric tests (classical and Bayesian paired *t*-tests using the Bayes Factor Toolbox^59^ with default priors, repeated-measures ANOVA). Tests were performed in a paired manner across participants and/or model predictions, using MATLAB built-in functions. Effect sizes are reported as Cohen’s d for paired samples and, for clusters, as the mean Cohen’s d across the samples of the cluster. For repeated-measures ANOVA, reported *p*-values are Greenhouse-Geisser corrected. Given our sample sizes, these statistical tests were applied outside the small-sample regime. Data were not explicitly tested for normality. Post-hoc statistics are corrected for multiple comparisons using the Tukey-Kramer method. Peak latencies and amplitudes, and the relative compression in ROIs, were estimated with a jackknife procedure (leave-one-participant-out) and compared with jackknife-corrected statistics^60,61^. Time windows for averaging were defined as the samples where the group-average decoding precision reached 90% of its peak. In multiple regressions, the unique contribution of each predictor was measured as the decrease in *r*^2^ when this predictor was removed. Inter-individual correlations were computed across participants at each time sample (Pearson correlation, with log-transformed inference noise and bootstrapped s.e.m.), and multiple regressions used z-scored variables. Given the absence of prior effect sizes, we chose a sample size (N = 37) which exceeded the average sample size used in human psychophysical studies with similar trial number per participant (n = 540). Cluster-based permutation testing^62^ was used to correct for multiple comparisons. Data points with *p* < 0.05 were grouped into same-sign clusters and scored by their summed |t|. Cluster-corrected p-values were obtained by comparing each cluster’s statistic to a null distribution of maximum cluster statistics from 10,000 permutations (condition permutation, subject permutation or RNN permutation).

## Acknowledgements

This work was supported by the European Research Council (ERC-StG 759341 to Valentin Wyart), the National Institute of Mental Health (US–France collaborative research grant No. 1R01MH115554-01 to Jan Drugowitsch and Valentin Wyart), the Agence Nationale de la Recherche (grant No. ANR-17-NEUC-0001-02 to Valentin Wyart and Jan Drugowitsch), a postdoctoral grant from the Fondation pour la Recherche Médicale (SPF-202309017516 to Lucas Benjamin) and a department-wide grant No. ANR-17-EURE-0017 (to the Département d’Études Cognitives of the École Normale Supérieure). The funders had no role in study design, data collection and analysis, decision to publish or preparation of the manuscript.

## Author contributions

Julie Drevet: Conceptualization, Data curation, Formal analysis, Investigation, Methodology, Software, Validation, Visualization, Writing – original draft, Writing – review & editing. Lucas Benjamin: Formal analysis, Methodology, Software, Writing – review & editing. Lindsay Rondot: Investigation, Writing – review & editing. Jan Drugowitsch: Conceptualization, Funding acquisition, Methodology, Writing – review & editing. Valentin Wyart: Conceptualization, Formal analysis, Funding acquisition, Methodology, Project administration, Resources, Software, Supervision, Visualization, Writing – original draft, Writing – review & editing.

## Data availability

The behavioral data are available with the analysis code. The preprocessed MEG data of each participant (continuous MEG data, events and source models) and the source-level decoding results are available on Zenodo at https://doi.org/10.5281/zenodo.23266592.

## Code availability

The code used to analyze the behavioral and MEG data, to fit the computational models, and to train and analyze the recurrent neural networks (MATLAB R2022b and Python 3.11) is available on GitHub at https://github.com/juliedrevet/NEURCOMP and archived on Zenodo: https://doi.org/10.5281/zenodo.23667406. The trained networks can be regenerated with the provided code. Claude (Anthropic) was used to help reorganize, document and check the analysis code for public release, and to proofread an earlier version of the manuscript. Julie Drevet verified the codebase and takes full responsibility for its contents.

## References

1. Gold, J. I. & Shadlen, M. N. The neural basis of decision making. Annu. Rev. Neurosci. 30, 535–574 (2007).

2. Wald, A. & Wolfowitz, J. Optimum character of the sequential probability ratio test. Ann. Math. Stat. 19, 326–339 (1948).

3. Bogacz, R., Brown, E., Moehlis, J., Holmes, P. & Cohen, J. D. The physics of optimal decision making: a formal analysis of models of performance in two-alternative forced-choice tasks. Psychol. Rev. 113, 700–765 (2006).

4. Rahnev, D. & Denison, R. N. Suboptimality in perceptual decision making. Behav. Brain Sci. 41, e223 (2018).

5. Wyart, V. & Koechlin, E. Choice variability and suboptimality in uncertain environments. Curr. Opin. Behav. Sci. 11, 109–115 (2016).

6. Duffy, J. S., Bellgrove, M. A., Murphy, P. R. & O’Connell, R. G. Disentangling sources of variability in decision-making. Nat. Rev. Neurosci. 26, 247–262 (2025).

7. Drugowitsch, J., Wyart, V.Devauchelle, A.-D. & Koechlin, E. Computational precision of mental inference as critical source of human choice suboptimality. Neuron 92, 1398–1411 (2016).

8. Denève, S., Latham, P. E. & Pouget, A. Efficient computation and cue integration with noisy population codes. Nat. Neurosci. 4, 826–831 (2001).

9. Jazayeri, M. & Movshon, J. A. Optimal representation of sensory information by neural populations. Nat. Neurosci. 9, 690–696 (2006).

10. Pouget, A., Dayan, P. & Zemel, R. S. Inference and computation with population codes. Annu. Rev. Neurosci. 26, 381–410 (2003).

11. Amari, S. Dynamics of pattern formation in lateral-inhibition type neural fields. Biol. Cybern. 27, 77–87 (1977).

12. Ermentrout, B. Neural networks as spatio-temporal pattern-forming systems. Rep. Prog. Phys. 61, 353–430 (1998).

13. Esnaola-Acebes, J. M., Roxin, A. & Wimmer, K. Flexible integration of continuous sensory evidence in perceptual estimation tasks. Proc. Natl Acad. Sci. USA 119, e2214441119 (2022).

14. Ma, W. J., Beck, J. M., Latham, P. E. & Pouget, A. Bayesian inference with probabilistic population codes. Nat. Neurosci. 9, 1432–1438 (2006).

15. van Bergen, R. S., Ma, W. J., Pratte, M. S. & Jehee, J. F. M. Sensory uncertainty decoded from visual cortex predicts behavior. Nat. Neurosci. 18, 1728–1730 (2015).

16. Li, H.-H., Sprague, T. C., Yoo, A. H., Ma, W. J. & Curtis, C. E. Joint representation of working memory and uncertainty in human cortex. Neuron 109, 3699–3712 (2021).

17. Geurts, L. S., Cooke, J. R. H., van Bergen, R. S. & Jehee, J. F. M. Subjective confidence reflects representation of Bayesian probability in cortex. Nat. Hum. Behav. 6, 294–305 (2022).

18. Shadlen, M. N., Kiani, R., Hanks, T. D. & Churchland, A. K. in Better Than Conscious? Decision Making, the Human Mind, and Implications for Institutions (eds Engel, C. & Singer, W.) 71–102 (MIT Press, 2008).

19. Shadlen, M. N. & Kiani, R. Decision making as a window on cognition. Neuron 80, 791–806 (2013).

20. Shushruth, S., Zylberberg, A. & Shadlen, M. N. Sequential sampling from memory underlies action selection during abstract decision-making. Curr. Biol. 32, 1949–1960 (2022).

21. O’Connell, R. G., Dockree, P. M. & Kelly, S. P. A supramodal accumulation-to-bound signal that determines perceptual decisions in humans. Nat. Neurosci. 15, 1729–1735 (2012).

22. Twomey, D. M., Kelly, S. P. & O’Connell, R. G. Abstract and effector-selective decision signals exhibit qualitatively distinct dynamics before delayed perceptual reports. J. Neurosci. 36, 7346–7352 (2016).

23. Freedman, D. J. & Assad, J. A. Experience-dependent representation of visual categories in parietal cortex. Nature 443, 85–88 (2006).

24. Wyart, V., de Gardelle, V., Scholl, J. & Summerfield, C. Rhythmic fluctuations in evidence accumulation during decision making in the human brain. Neuron 76, 847–858 (2012).

25. Barlow, H. B. in Sensory Communication (ed. Rosenblith, W. A.) 217–234 (MIT Press, 1961).

26. Usher, M. & McClelland, J. L. The time course of perceptual choice: the leaky, competing accumulator model. Psychol. Rev. 108, 550–592 (2001).

27. Kiani, R., Hanks, T. D. & Shadlen, M. N. Bounded integration in parietal cortex underlies decisions even when viewing duration is dictated by the environment. J. Neurosci. 28, 3017–3029 (2008).

28. Ossmy, O. et al. The timescale of perceptual evidence integration can be adapted to the environment. Curr. Biol. 23, 981–986 (2013).

29. Weiss, A., Chambon, V., Lee, J. K., Drugowitsch, J. & Wyart, V. Interacting with volatile environments stabilizes hidden-state inference and its brain signatures. Nat. Commun. 12, 2228 (2021).

30. Salvador, A. et al. Premature commitment to uncertain decisions during human NMDA receptor hypofunction. Nat. Commun. 13, 338 (2022).

31. Drevet, J., Drugowitsch, J. & Wyart, V. Efficient stabilization of imprecise statistical inference through conditional belief updating. Nat. Hum. Behav. 6, 1691–1704 (2022).

32. Palminteri, S., Wyart, V. & Koechlin, E. The importance of falsification in computational cognitive modeling. Trends Cogn. Sci. 21, 425–433 (2017).

33. Wilson, R. C. & Collins, A. G. E. Ten simple rules for the computational modeling of behavioral data. eLife 8, e49547 (2019).

34. Polanía, R., Woodford, M. & Ruff, C. C. Efficient coding of subjective value. Nat. Neurosci. 22, 134–142 (2019).

35. Finkelstein, A. et al. Attractor dynamics gate cortical information flow during decision-making. Nat. Neurosci. 24, 843–850 (2021).

36. Tsetsos, K. et al. Economic irrationality is optimal during noisy decision making. Proc. Natl Acad. Sci. USA 113, 3102–3107 (2016).

37. Li, V., Herce Castañón, S., Solomon, J. A., Vandormael, H. & Summerfield, C. Robust averaging protects decisions from noise in neural computations. PLoS Comput. Biol. 13, e1005723 (2017).

38. Spitzer, B., Waschke, L. & Summerfield, C. Selective overweighting of larger magnitudes during noisy numerical comparison. Nat. Hum. Behav. 1, 0145 (2017).

39. Freedman, D. J., Riesenhuber, M., Poggio, T. & Miller, E. K. Categorical representation of visual stimuli in the primate prefrontal cortex. Science 291, 312–316 (2001).

40. Bennur, S. & Gold, J. I. Distinct representations of a perceptual decision and the associated oculomotor plan in the monkey lateral intraparietal area. J. Neurosci. 31, 913–921 (2011).

41. Freedman, D. J. & Assad, J. A. A proposed common neural mechanism for categorization and perceptual decisions. Nat. Neurosci. 14, 143–146 (2011).

42. Mante, V., Sussillo, D., Shenoy, K. V. & Newsome, W. T. Context-dependent computation by recurrent dynamics in prefrontal cortex. Nature 503, 78–84 (2013).

43. Pagan, M. et al. Individual variability of neural computations underlying flexible decisions. Nature 639, 421–429 (2025).

44. Benjamin, L., Drevet, J. & Wyart, V. Neural abstraction of evidence accumulation enables flexible multidimensional perceptual decisions. Preprint at 10.64898/2026.10.07.757231 (2026).

45. Flesch, T., Juechems, K., Dumbalska, T., Saxe, A. & Summerfield, C. Orthogonal representations for robust context-dependent task performance in brains and neural networks. Neuron 110, 1258–1270 (2022).

46. van Ede, F. & Nobre, A. C. Turning attention inside out: how working memory serves behavior. Annu. Rev. Psychol. 74, 137–165 (2023).

47. Findling, C. & Wyart, V. Computation noise promotes zero-shot adaptation to uncertainty during decision-making in artificial neural networks. Sci. Adv. 10, eadl3931 (2024).

48. Ji-An, L., Benna, M. K. & Mattar, M. G. Discovering cognitive strategies with tiny recurrent neural networks. Nature 644, 993–1001 (2025).

49. Eckstein, M. K., Summerfield, C., Daw, N. D. & Miller, K. J. Hybrid neural–cognitive models reveal how memory shapes human reward learning. Nat. Hum. Behav. 10, 972–987 (2026).

50. Molano-Mazón, M. et al. Recurrent networks endowed with structural priors explain suboptimal animal behavior. Curr. Biol. 33, 622–638 (2023).

51. Knowlton, B. J., Mangels, J. A. & Squire, L. R. A neostriatal habit learning system in humans. Science 273, 1399–1402 (1996).

52. Acerbi, L. Variational Bayesian Monte Carlo. in Advances in Neural Information Processing Systems vol. 31, 8213–8223 (Curran Associates, Inc., 2018).

53. Acerbi, L. Variational Bayesian Monte Carlo with noisy likelihoods. in Advances in Neural Information Processing Systems vol. 33, 8211–8222 (Curran Associates, Inc., 2020).

54. Oostenveld, R., Fries, P., Maris, E. & Schoffelen, J.-M. FieldTrip: open source software for advanced analysis of MEG, EEG, and invasive electrophysiological data. Comput. Intell. Neurosci. 2011, 156869 (2011).

55. Fischl, B. FreeSurfer. NeuroImage 62, 774–781 (2012).

56. Tadel, F., Baillet, S., Mosher, J. C., Pantazis, D. & Leahy, R. M. Brainstorm: a user-friendly application for MEG/EEG analysis. Comput. Intell. Neurosci. 2011, 879716 (2011).

57. Glorot, X. & Bengio, Y. Understanding the difficulty of training deep feedforward neural networks. in Proceedings of the Thirteenth International Conference on Artificial Intelligence and Statistics vol. 9, 249–256 (JMLR Workshop and Conference Proceedings, 2010).

58. Saxe, A. M., McClelland, J. L. & Ganguli, S. Exact solutions to the nonlinear dynamics of learning in deep linear neural networks. Preprint at https://arxiv.org/abs/1312.6120 (2014).

59. Krekelberg, B. Matlab Toolbox for Bayes Factor Analysis. Zenodo 10.5281/zenodo.4394422 (2024).

60. Miller, J., Patterson, T. & Ulrich, R. Jackknife-based method for measuring LRP onset latency differences. Psychophysiology 35, 99–115 (1998).

61. Ulrich, R. & Miller, J. Using the jackknife-based scoring method for measuring LRP onset effects in factorial designs.Psychophysiology 38, 816–827 (2001).

62. Maris, E. & Oostenveld, R. Nonparametric statistical testing of EEG- and MEG-data. J. Neurosci. Methods 164, 177–190 (2007).

