## Supplementary Information for "Flexible compression of sensory evidence makes human decisions more resilient to noisy neural computations"

### Supplementary Figures

**a** generative distributions on one trial

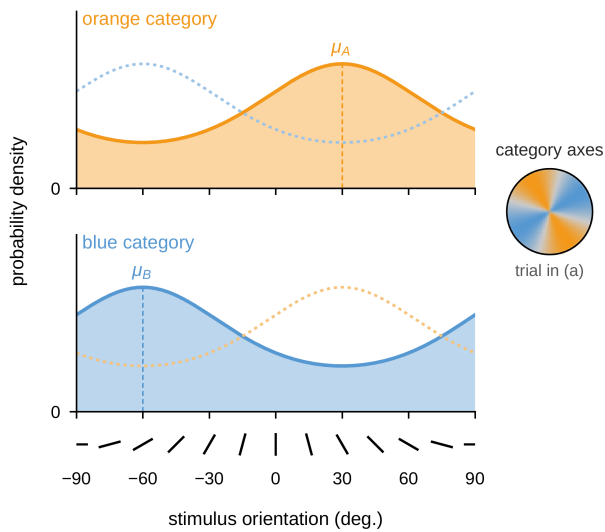

**b** evidence provided by each stimulus across trials

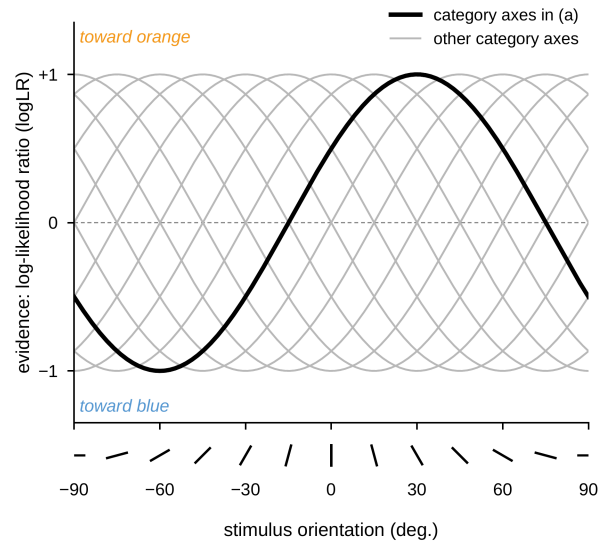

**Supplementary Figure 1 | Generative distributions and evidence provided by each stimulus.** **(a)** Stimuli correspond to bars of orientations $\theta_1, \dots, \theta_n$ drawn from one of two color-labeled categories, associated with overlapping von Mises probability density functions (concentration $\kappa = 0.5$) centered on orthogonal orientations, the category axes $\mu_A$ (orange) and $\mu_B$ (blue). Top: Orange category. Bottom: Blue category. In each panel, the probability density function of the other category is shown as a light dotted line. In this example sequence, the category axes are at $+30^\circ$ (orange) and $-60^\circ$ (blue), as depicted by the inset. Oriented bars below the x-axis illustrate stimulus orientations ($0^\circ$: vertical). **(b)** Evidence provided by each stimulus orientation $\theta$ in favor of the orange category, expressed as its log-likelihood ratio $\log LR = 2\kappa \cos(2(\theta - \mu_A))$. The black line corresponds to the category axes shown in (a), and gray lines to other positions of the category axes (in steps of $15^\circ$). Because the category axes changed unpredictably from one sequence to the next, the same stimulus orientation provided evidence of different strength and sign across sequences.

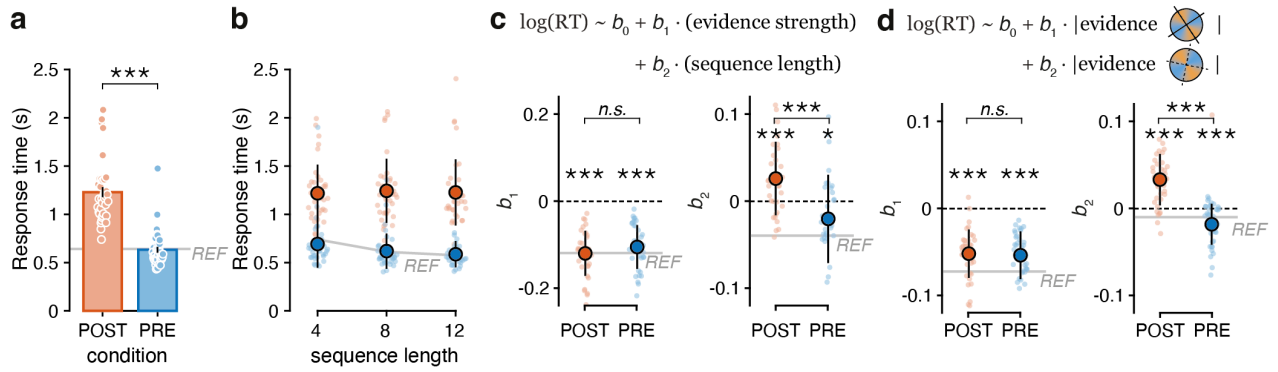

**Supplementary Figure 2 | Effects of accumulation space on response times (N = 37).** **(a)** Mean response times measured from 'go'-signal, in each condition. Participants take overall more time to report their choice in POST, but are similarly fast in PRE and REF. Bars and error bars indicate mean ± s.e.m., small dots indicate single participants. **(b)** Response times depending on sequence length for each condition. No effect of sequence length on response times found in POST, but participants are faster for longer sequences in PRE and REF conditions. Color dots with error bars indicate mean and s.d., small color dots indicate single participants. **(c)** Predicted effect of evidence strength and sequence length on response times. Left: Evidence strength regression coefficient $b_1$. Stronger sequence evidence decreases response times similarly in all conditions. Right: Sequence length regression coefficient $b_2$. Opposite effect of sequence length depending on condition. Color dots with error bars indicate mean and s.d., small color dots indicate single participants. **(d)** Predicted effect of absolute evidence strength relative to true and null projection axes. Left: True evidence strength regression coefficient $b_1$. Relevant information speeds up the response times similarly in both POST and PRE. Right: Null evidence strength regression coefficient $b_2$. Irrelevant information slows down participants in POST but not in the other two conditions. Color dots with error bars indicate mean and s.d., small color dots indicate single participants. In all panels: the solid gray lines correspond to participants' behavior in REF, three stars and one star indicate a significant effect at $p < 0.001$ and $p < 0.05$, respectively, n.s. indicates no statistical difference.

**a** evidence accumulation spaces

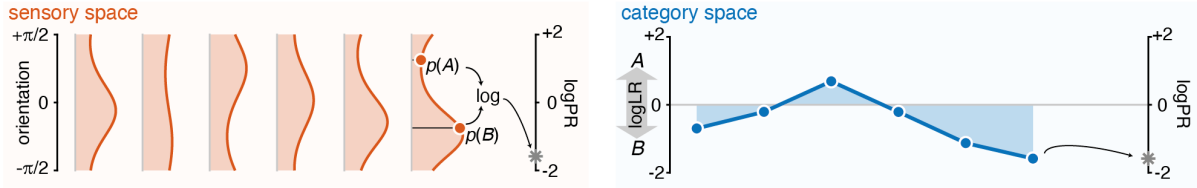

**b** best-fitting model parameters depending on accumulation space

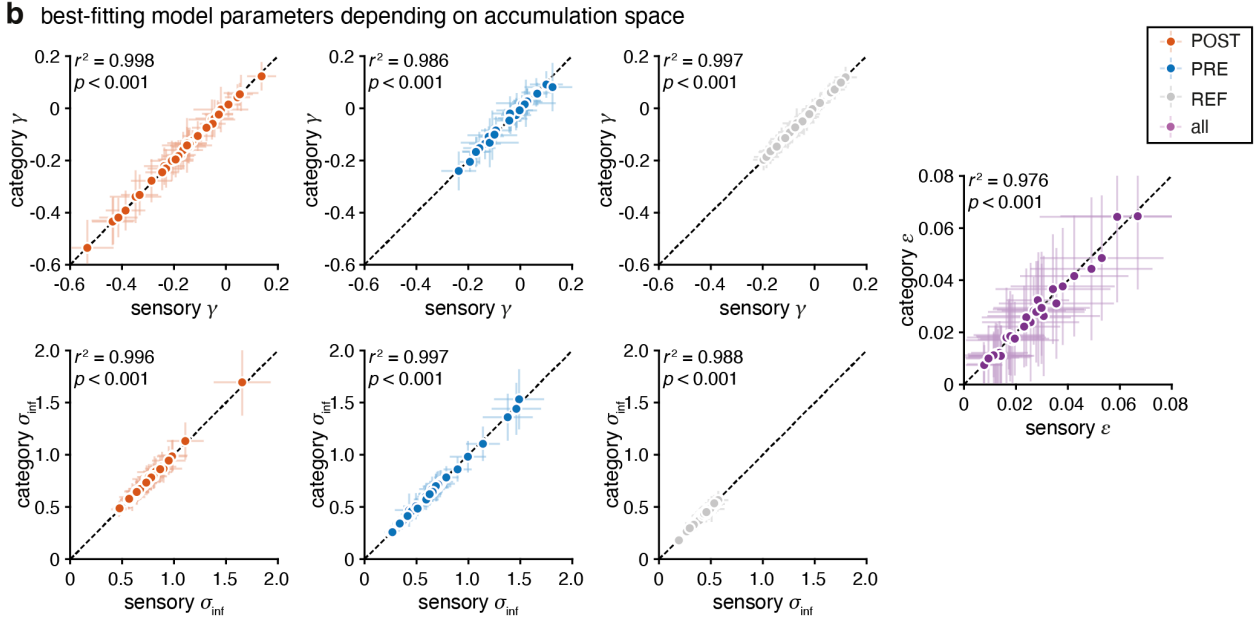

**Supplementary Figure 3 | Equivalence in best-fitting parameters of sensory and category space accumulation models**

**(N = 37).** **(a)** Evidence accumulation spaces (same as Fig. 1a). **(b)** In each condition and for each suboptimality parameter (inference imbalance $\gamma$, inference noise $\sigma_{\text{inf}}$ and lapse parameter $\epsilon$) the best-fitting value of the parameter from the computational model used in the main text (the accumulated variable is the evidence already categorized as log-likelihood ratio) was correlated to the best-fitting value of the parameter from the model accumulating evidence in sensory space (the accumulated variable is the full posterior). All parameters are almost perfectly correlated (all $r^2 > 0.9$, $p < 0.001$). Dots and error bars correspond to mean and s.d. of the best-fitting parameter for each human participant.

**a** model and parameter recovery on simulated data

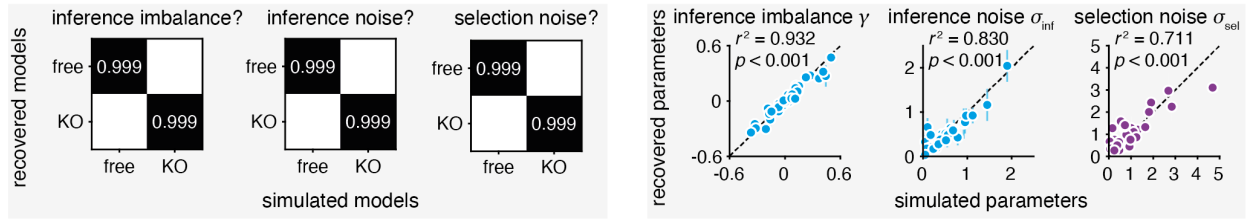

**b** best-fitting lapse parameter

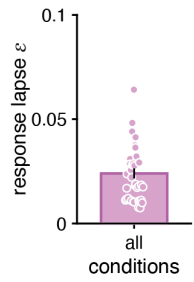

**c** correlation between best-fitting parameters

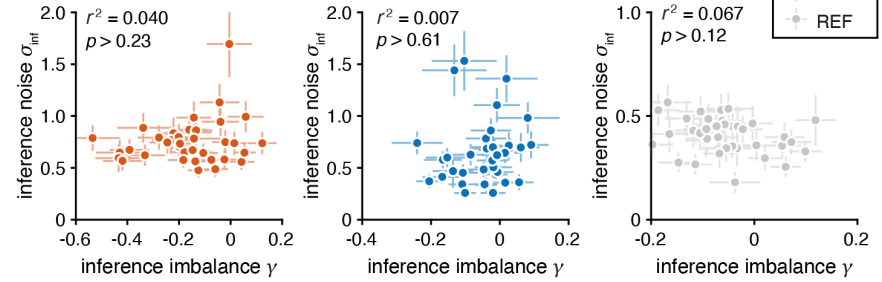

**Supplementary Figure 4 | Computational model. (a)** Model and parameter recovery. Left: Confusion matrices between model variants enabling or disabling each source of suboptimality depicting exceedance probabilities obtained when recovering simulated data. The presence and absence of each parameter in the ‘ground-truth’ model variants used to simulate synthetic behavior was correctly recovered with exceedance probabilities above 0.999. Right: Parameter recovery: the values of each source of suboptimal choices were adequately recovered as exhibited by the high correlation between simulated and recovered parameter values (squared Pearson correlation coefficient for the inference imbalance $\gamma$: $r^2 = 0.932$; for the inference noise $\sigma_{\text{inf}}$: $r^2 = 0.830$; for the selection noise $\sigma_{\text{sel}}$: $r^2 = 0.711$). Dots and error bars correspond to mean and s.d. **(b)** Best-fitting lapse parameter. A lapse parameter $\epsilon$ common to all conditions was fitted together with the other parameters and resulted in a low proportion of random responses ($2.42 \pm 0.24\%$, mean ± s.e.m.). Bar and error bar correspond to mean ± s.e.m., dots correspond to single participants. **(c)** Correlation between best-fitting model parameters inside each condition. Inference imbalance $\gamma$ and inference noise $\sigma_{\text{inf}}$ are not correlated. Dots and error bars correspond to mean and s.d.

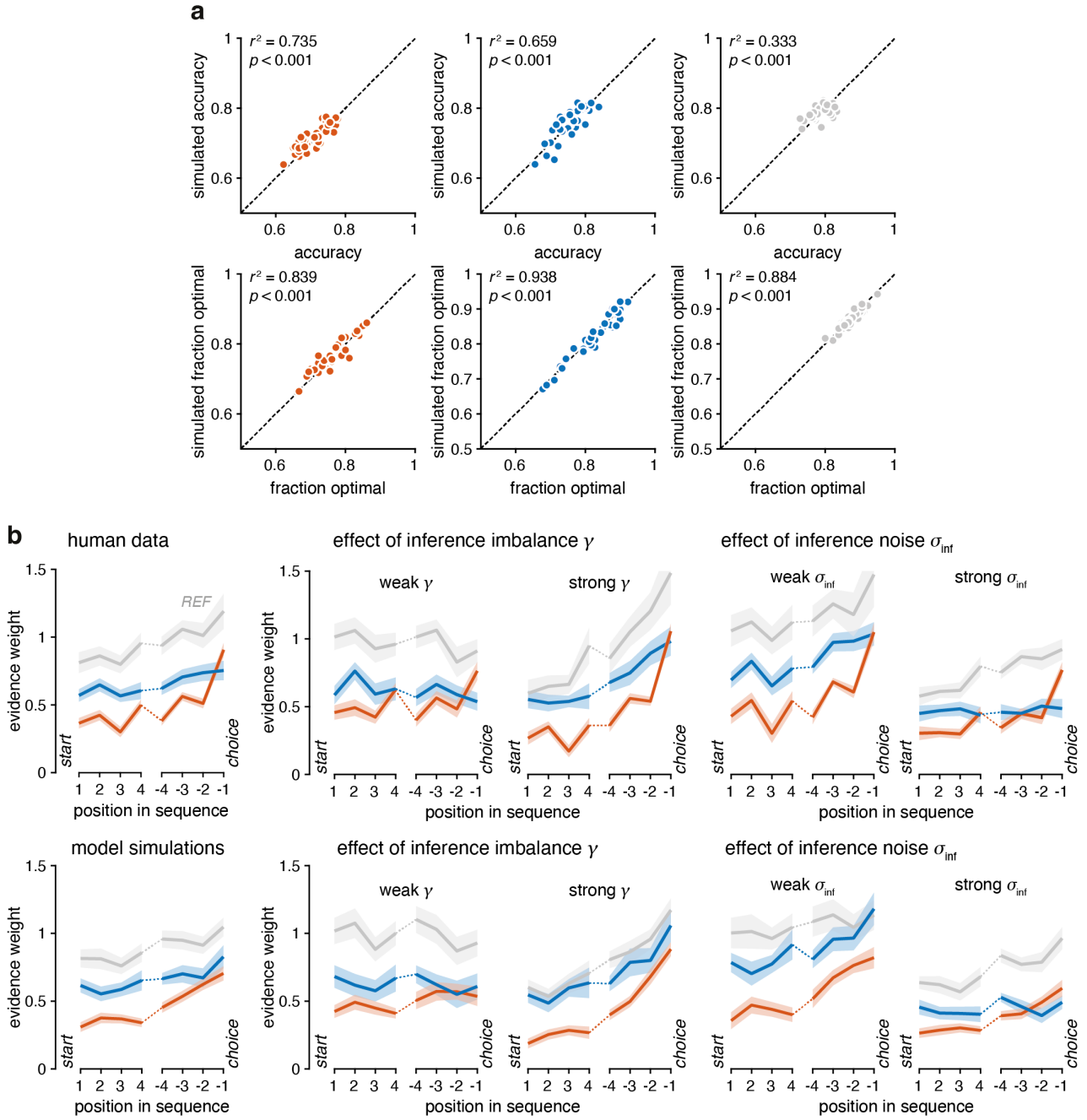

**Supplementary Figure 5 | Best-fitting model simulations successfully capturing human behavior (N = 37).** **(a)** Correlation between simulated and human accuracy (top) or simulated and human fraction of optimal choices (bottom) in each condition. For each participant, the model was simulated using the participant's best-fitting $\gamma$, $\sigma_{inf}$ and $\epsilon$ parameters and evaluated on the same experimental sequences as the participant. Dots indicate single participants. **(b)** Distinct gradients of behavioral variability associated with the inference imbalance and the inference noise parameters. The integration kernel (as seen in main Fig. 2d) for human participants (top row) or model simulations (lower row) obtained by sorting participants in two median-split groups as a function of the best-fitting value of each parameter (left: no sorting; middle: median-split on inference imbalance; right: median-split on inference noise). Model simulations qualitatively capture the behavioral variability of participants. Strong inference imbalance (i.e., strongly negative values, corresponding to a leak) results in a pronounced recency effect, whereas strong inference noise decreases the overall weight given to each piece of evidence. Lines and shaded areas correspond to mean ± s.e.m.

**a** neural coding of stimulus orientation for correct and error trials

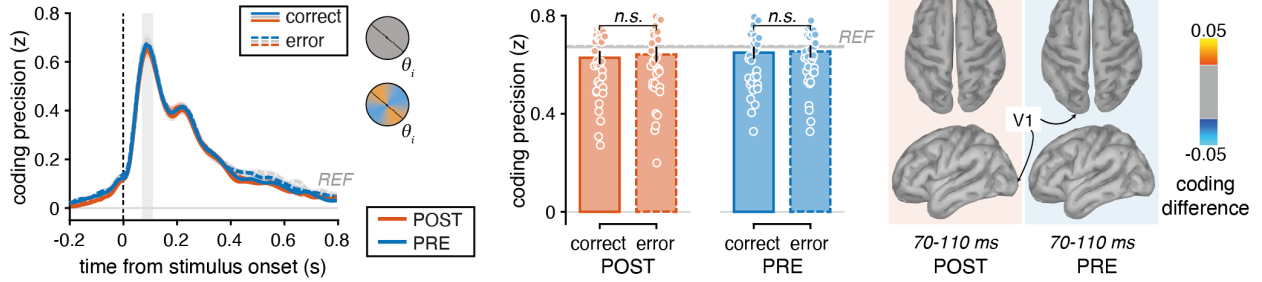

**b** neural coding of evidence strength for correct and error trials

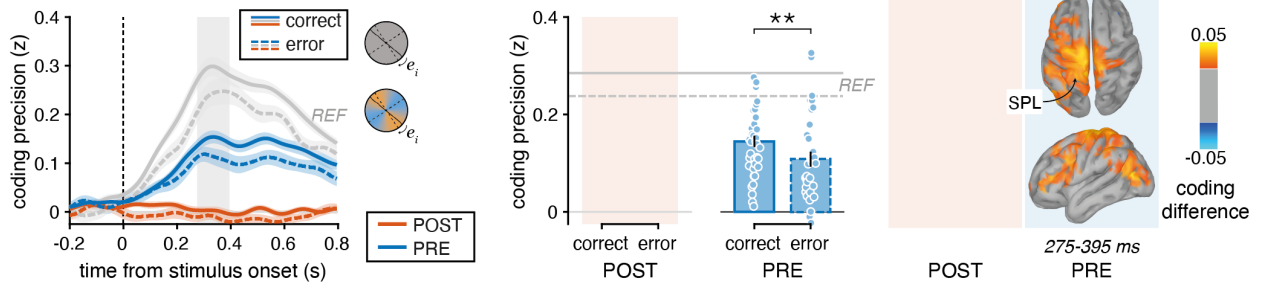

**Supplementary Figure 6 | Neural coding depending on choice accuracy (N = 37).** **(a)** Neural coding of stimulus orientation for correct and error trials. Left: Time course of neural coding for correct (solid) and error (dashed) trials. The gray shaded area corresponds to the [+70 +110] ms time window, during which the coding precision (irrespective of choice accuracy) reaches 90% of its peak value across conditions. Middle: Comparison of coding precision between correct and error trials. Coding is averaged on the time windows defined by the gray shaded area in the left panel. No difference in coding of stimulus orientation between correct and error trials. Bars and error bars correspond to mean ± s.e.m. and dots correspond to single participants. Right: Difference in coding precision between correct and error trials across the cortical surface in POST (left) and PRE (right) when averaged on the [+70 +110] ms time window. Difference thresholded at 50% of the maximum absolute difference. **(b)** Neural coding of evidence strength for correct and error trials. Left: Time course of neural coding for correct (solid) and error (dashed) trials. The gray shaded area corresponds to the [+275 +395] ms time window, during which the coding precision (irrespective of choice accuracy) reaches 90% of its peak value in PRE (first peak). Middle: Comparison of coding precision between correct and error trials. Coding is averaged on the condition-specific time windows during which the coding precision reaches 90% of its peak value (PRE: [+275 +395] ms, gray shaded area from the left panel; REF: [+280 +445] ms). Evidence strength is not decoded in POST; in PRE and REF evidence strength coding precision is significantly higher for correct than for error trials. Bars and error bars correspond to mean ± s.e.m. and dots correspond to single participants. Right: Difference in coding precision between correct and error trials across the cortical surface in POST (left, not decoded) and PRE (right) when averaged on the [+275 +395] ms window. Difference thresholded at 50% of the maximum absolute difference. In all panels: lines and shaded areas correspond to mean ± s.e.m., two stars indicate a significant effect at $p < 0.01$ and n.s. indicates no statistical difference.

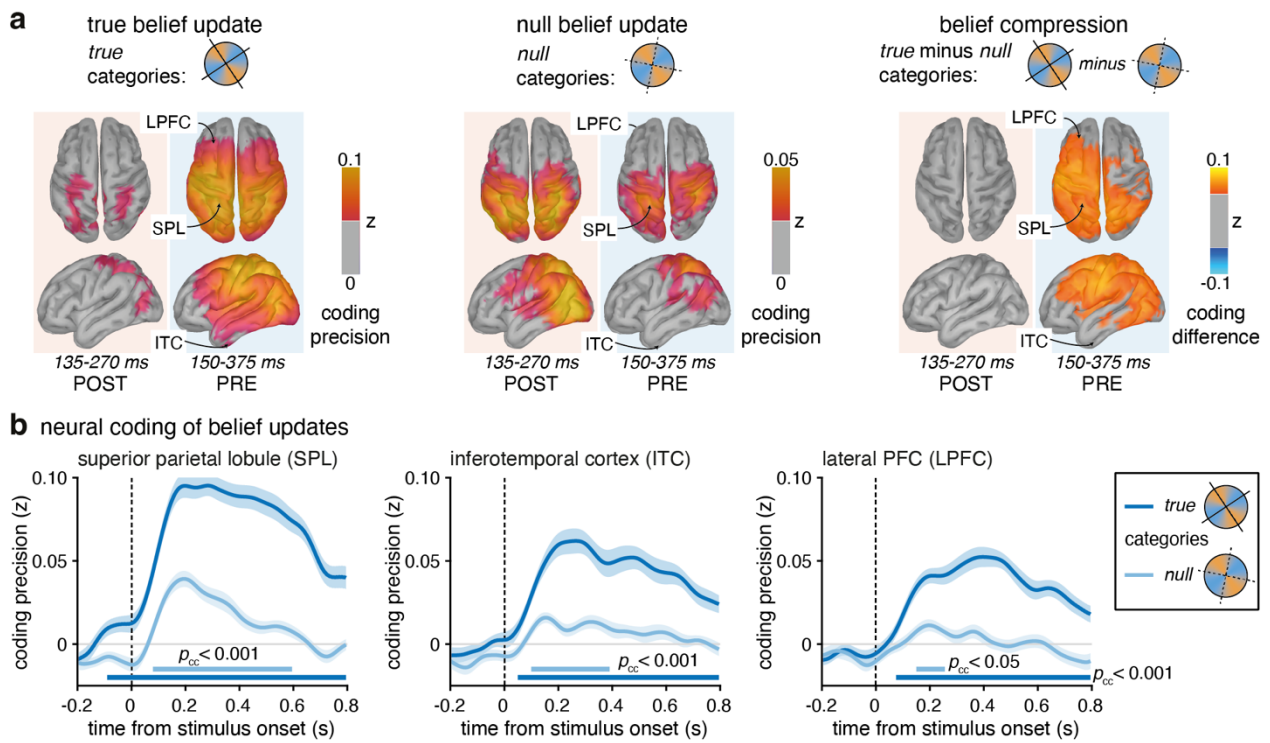

**Supplementary Figure 7 | Neural coding of belief updates across cortical surface (N = 37).** **(a)** Searchlight-based coding of belief update depending on category axis. Left: Belief update according to true categories. Coding is averaged on the condition-specific time windows during which the coding precision reaches 90% of its peak value at the sensor level (POST: [+135 +270] ms; PRE: [+150 +375] ms). The coding is thresholded at 50% of the coding peak from PRE (actual maximum coding precision: 0.1148). Middle: Belief update according to null categories. Coding is averaged on the same time window as for the true belief update (left panel). The coding is thresholded at 50% of the coding peak from POST (actual maximum coding precision: 0.0604). Right: Compression on decision-relevant axis as difference in coding precision between true and null belief update across the cortical surface. Difference thresholded at 50% of the maximum absolute difference (actual maximum coding difference: 0.0783). **(b)** Time course of the neural coding of belief update for three regions of interest (left: superior parietal lobule; middle: inferotemporal cortex; right: lateral prefrontal cortex) computed with respect to the true categories (dark blue) and the null categories (light blue). The true belief update is significantly decoded in all three regions (cluster-corrected $p < 0.001$), while the null belief update is significantly represented in the SPL and the ITC (cluster-corrected $p < 0.001$) but only weakly and briefly in the LPFC activity (cluster-corrected $p < 0.05$). Solid lines correspond to mean coding precision and shaded areas correspond to s.e.m.

**a** neural coding of true belief update for correct and error trials

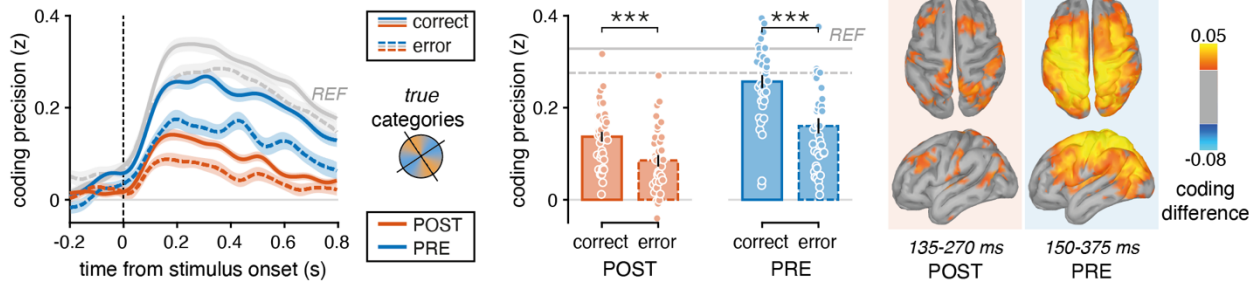

**b** neural coding of null belief update for correct and error trials

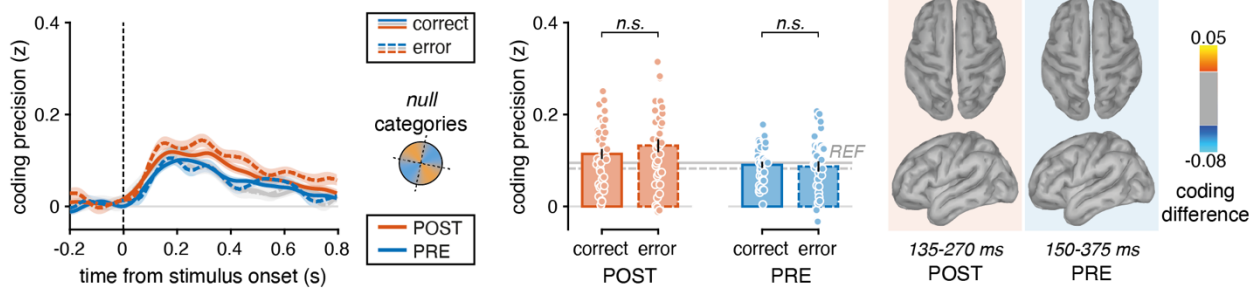

**Supplementary Figure 8 | Neural coding of belief update depending on category axis for correct and error trials (N = 37).**

**(a)** Neural coding of belief update computed with respect to the true categories for correct and error trials. Left: Time course for correct (solid) and error (dashed) trials. Middle: Comparison of coding precision between correct and error trials. Coding is averaged on the condition-specific time windows during which the coding precision reaches 90% of its peak value (POST: [+135 +270] ms; PRE: [+150 +375] ms; REF: [+155 +415] ms). Right: Difference in coding precision between correct and error trials across the cortical surface in POST (left) and PRE (right) when averaged on the condition-specific time window. Difference thresholded at 50% of the maximum absolute difference. **(b)** Neural coding of belief update computed with respect to the null categories for correct and error trials. Left: Time course for correct (solid) and error (dashed) trials. Middle: Comparison of coding precision between correct and error trials. Coding is averaged on the condition-specific time windows computed for the true belief as specified in (a). Right: Difference in coding precision between correct and error trials across the cortical surface in POST (left) and PRE (right) when averaged on the condition-specific time window. Difference thresholded as in (a). In all panels: solid lines correspond to mean coding precision and shaded areas correspond to s.e.m., bars and error bars indicate mean ± s.e.m., small dots indicate single participants, the gray lines correspond to REF, three stars indicate a significant effect at $p < 0.001$ and n.s. indicates no statistical difference for one-tailed t-test testing correct > error.

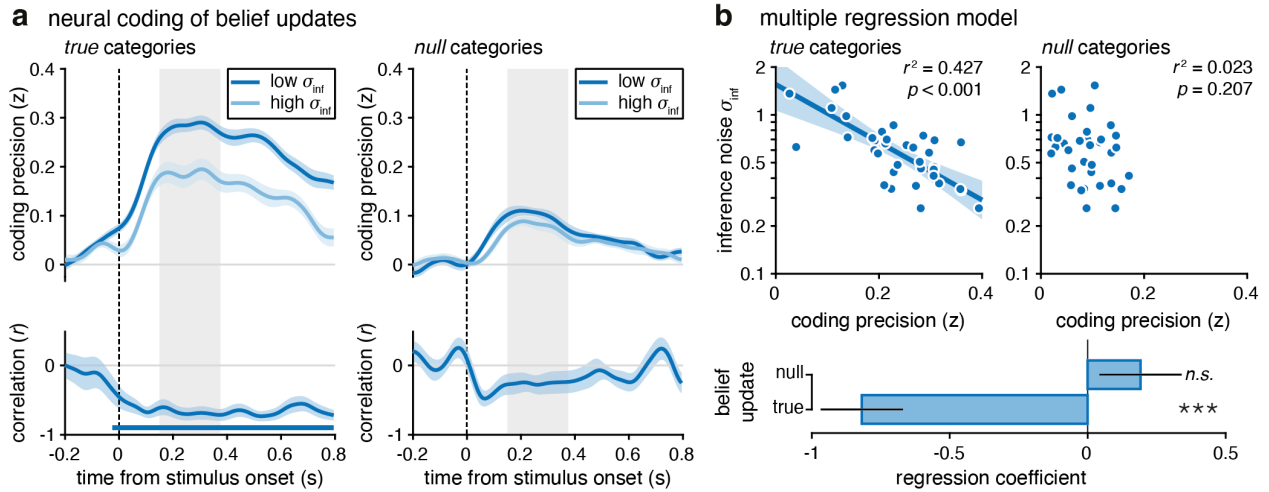

**Supplementary Figure 9 | Relation between inference noise and neural coding of belief updates in PRE (N = 37).** **(a)** Gradients in neural coding of the true belief update (left) and the null belief update (right) associated with inference noise. Top: Time course of neural coding when sorting participants in two median-split groups as a function of their best-fitting inference noise parameter $\sigma_{\text{inf}}$. Dark blue lines correspond to low values of $\sigma_{\text{inf}}$ (i.e., more precise inference) and light blue lines correspond to high values of $\sigma_{\text{inf}}$ (i.e., less precise inference). The gray shaded areas indicate the [+150 +375] ms time window, during which the coding precision reaches 90% of its peak value for the true belief update. Bottom: Time-resolved correlation between log-transformed $\sigma_{\text{inf}}$ and the belief updates. The true belief update correlates negatively with $\sigma_{\text{inf}}$ on the whole post-stimulus time window (cluster-corrected $p < 0.001$). No significant cluster for the correlation between null belief update and $\sigma_{\text{inf}}$. Lines and shaded areas correspond to mean ± s.e.m. Horizontal solid blue line indicates cluster of significance. **(b)** Multiple regression model predicting inference noise (log-transformed) parameter from true and null belief updates decodability. Decodability of each belief update corresponds to the mean neural coding on the [+150 +375] ms time window indicated by the gray shaded areas in (a). Top: Partial regression for true (left) and null (right) belief updates. Dots indicate single participants, lines indicate partial regression controlling for the alternative categories, shaded areas represent the 95% confidence interval. Bottom: Regression coefficients. Participants' inference noise parameter is strongly predicted by how participants represent the true belief update ($\beta = -0.818$, $p < 0.001$) but unrelated to how they represent the null belief update ($\beta = 0.191$, $p = 0.207$). Bars and error bars indicate coefficient ± s.e. Three stars indicate a significant effect at $p < 0.001$ and n.s. indicates no statistical difference.

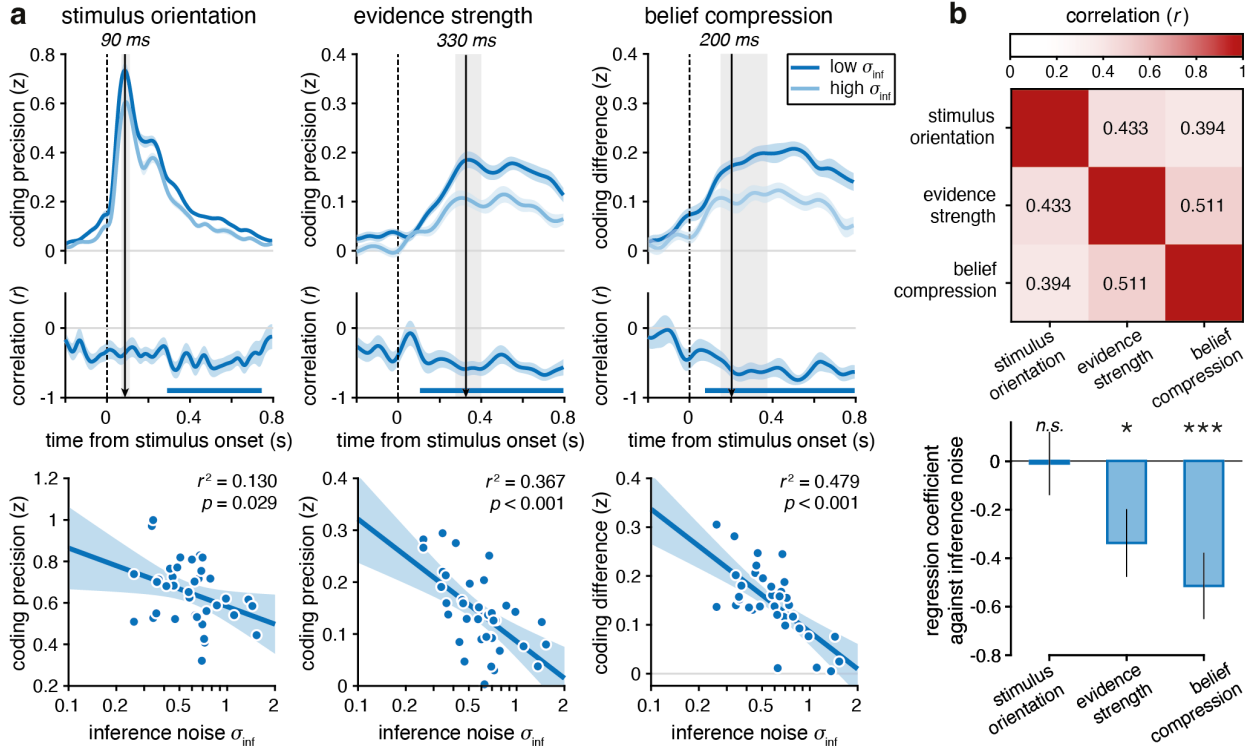

**Supplementary Figure 10 | Relation between inference noise and neural coding of stimulus characteristics in PRE (N = 37).** **(a)** Gradients in neural coding of stimulus orientation (left), evidence strength (middle) and belief compression (right) associated with inference noise. Top: Time course of neural coding when sorting participants in two median-split groups as a function of their best-fitting inference noise parameter $\sigma_{inf}$. Dark blue lines correspond to low values of $\sigma_{inf}$ (i.e., more precise inference) and light blue lines correspond to high values of $\sigma_{inf}$ (i.e., less precise inference). Vertical arrows indicate jackknifed mean latency of the peak value (for belief compression, the peak of the true belief update). The gray shaded areas indicate the time window, during which the coding precision reaches 90% of its peak value in PRE (stimulus orientation: [+70 +110] ms; evidence strength: [+275 +395] ms; belief compression evaluated for true belief update: [+150 +375] ms). Shaded areas correspond to s.e.m. Middle: Time-resolved correlation between log-transformed $\sigma_{inf}$ and the stimulus characteristic neural coding. Horizontal solid blue lines indicate clusters of significance (cluster-corrected $p < 0.001$). Bottom: Pearson correlation between log-transformed $\sigma_{inf}$ and the stimulus characteristic decodability (reported squared Pearson $r$). Decodability of each characteristic corresponds to the mean neural coding on the time window indicated by the gray shaded areas in the top row. Dots correspond to single participants, solid lines and shaded areas correspond to regression lines and 95% CI. **(b)** Assessing the unique relationship between belief compression and inference noise. Top: Correlation matrix between each stimulus characteristic decodability. Numbers indicate Pearson correlation coefficients $r$. Decodability of each characteristic corresponds to the mean neural coding on the time window indicated by the gray shaded areas in (a). Bottom: Multiple regression model coefficients. A higher belief compression is associated with lower inference noise even when accounting for the influence of stimulus orientation and evidence strength. Bars and error bars indicate coefficient ± s.e. Three stars and one star indicate a significant effect at $p < 0.001$ and $p < 0.05$, respectively, n.s. indicates no statistical difference.

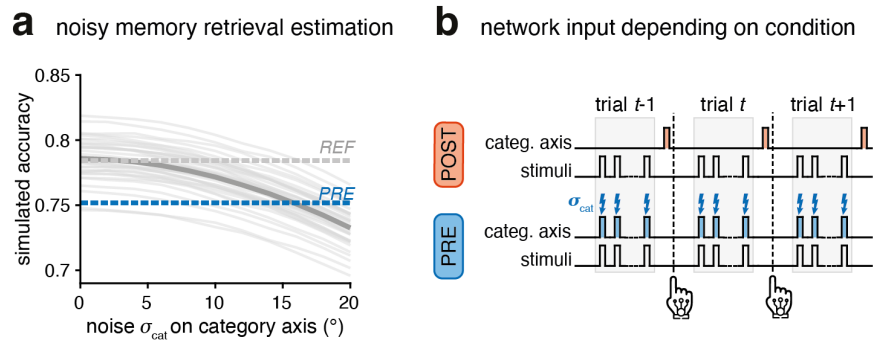

**Supplementary Figure 11 | Recurrent neural network specification. (a)** Noisy memory retrieval estimation. Decision accuracy simulated with the Bayesian suboptimal model, using inference imbalance $\gamma$ and inference noise $\sigma_{inf}$ fixed to their best-fitting values in the REF condition (categories visible throughout), with added angular noise $\sigma_{cat}$ on the category axis to capture imprecision from noisy memory retrieval. Participants' PRE accuracy is matched by approximately $\sigma_{cat} = 15^\circ$ (s.d.) of category-axis noise, providing a qualitative estimate of the imprecision with which decision categories are maintained in working memory. RNNs were provided with $\sigma_{cat} = 15^\circ$ noise on category axis in PRE sequences, approximating continuous noisy memory retrieval during training. Thin lines correspond to individual model simulations, the thick gray line corresponds to the mean simulated accuracy, and dotted lines correspond to participants' accuracy in the REF (gray) and PRE (blue) conditions. **(b)** Network input depending on condition. In the POST condition, the exact category axis is provided to the network at the end of the sequence. In the PRE condition, a noisy memory retrieval of the category axis is provided together with each stimulus orientation.

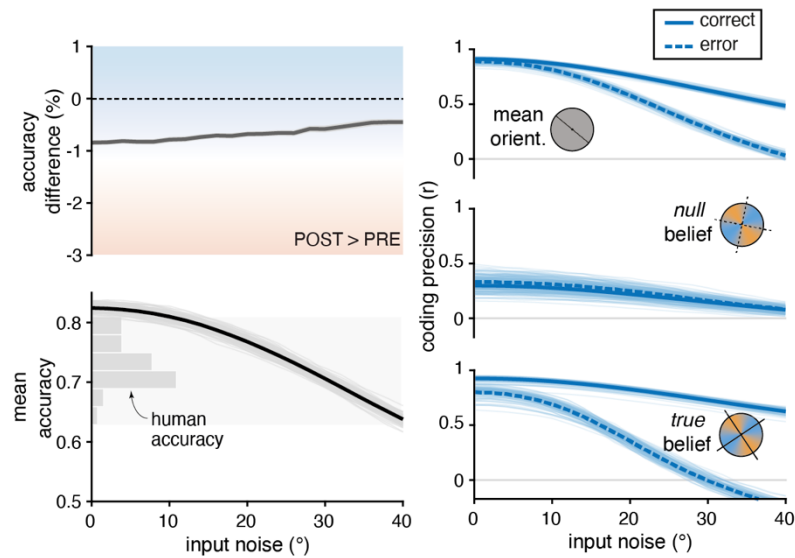

**Supplementary Figure 12 | Input noise does not produce a PRE accuracy advantage.** Same analyses as Fig. 7c, but injecting noise into the network input rather than the recurrent layer. Input noise was implemented by adding independent Gaussian noise to the orientation of each stimulus in the sequence, leaving recurrent dynamics unperturbed. Left: Accuracy as a function of input noise for PRE and POST trials. Top: Difference in accuracy between PRE and POST trials. Bottom: Mean accuracy across PRE and POST. Input noise reduced accuracy in both conditions, with no accuracy advantage for PRE over POST at any noise level matching participants' accuracy. Human PRE and POST mean accuracy is indicated by the gray histogram. Shaded areas indicate s.e.m. across RNNs. Right: Effect of input noise on the coding precision of the mean orientation of the sequence (top), null belief (middle) and true belief (bottom) for correct (solid lines) and error (dashed lines) trials from the PRE condition. Both mean orientation and true belief coding precision differed between correct and error trials, as under recurrent noise (Fig. 7c), but the difference was larger for the mean orientation (mean correct-error difference: $\Delta r = 0.19$, compared to 0.08 under recurrent noise), consistent with input noise degrading sensory evidence upstream of compression. Thin lines indicate individual RNNs.

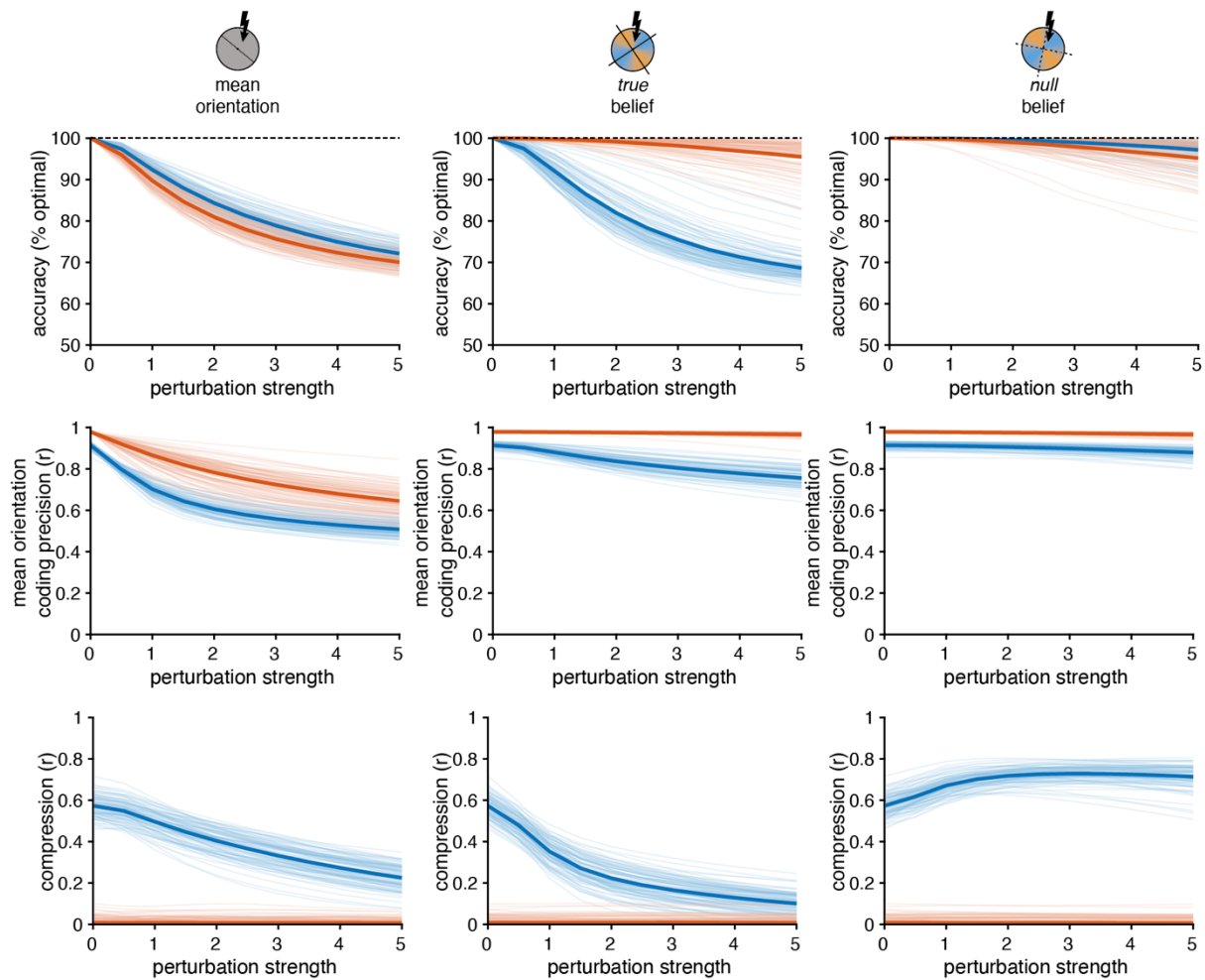

**Supplementary Figure 13 | Effect of causal perturbations.** Effect of perturbing the coding of mean orientation (left), true belief (middle), or null belief (right) in the recurrent layer, for PRE (blue) and POST (red) trials. Top: Accuracy (normalized to optimal performance without perturbation, dotted black line) as a function of perturbation strength. Perturbing mean orientation impairs POST more than PRE; perturbing true belief impairs PRE while leaving POST largely unaffected; perturbing null belief has negligible effect on either condition. Middle: Coding precision of mean orientation as a function of perturbation strength, shown separately for each perturbation type. Mean orientation coding is degraded by directly perturbing its own representation, but comparatively spared when perturbing true or null belief instead, confirming the specificity of each perturbation. Bottom: Neural compression (coding precision of true minus null beliefs) as a function of perturbation strength. Compression decreased when perturbing mean orientation or true belief in PRE, but increased when perturbing null belief, and remained near zero in POST throughout. Thin lines indicate individual RNNs, thick lines indicate mean across RNNs.

### Supplementary Results

#### Effect on response times

We examined the effect of condition on another behavioral signature of perceptual decision-making: response times (**Supplementary Fig. 2**). Due to the task design, in POST, participants were presented with the category-mapping cue together with the ‘go’ signal at the end of the sequence. Therefore, we expected participants to be significantly slower than in PRE because of the time needed to first map the inferred mean of the sequence to these categories. We indeed found clearly delayed response times in POST, but still correlated with response times in PRE (difference POST-PRE on log(RT): $t_{36} = 25.8$, $p < 0.001$; linear correlation coefficient $r = 0.777$, $p < 0.001$, 95% CI = [0.60 0.88]; **Supplementary Fig. 2a**). However, in PRE, when participants could relate each stimulus to its underlying category, they were not significantly slower to respond than in REF (post-hoc $t$-test, difference REF-PRE on log(RT): $t_{36} = 1.57$, $p = 0.27$, $BF_{01} = 1.84$). Interestingly, while accuracy increased with sequence length in all conditions (all $F_{2,72} > 13.4$, all $p < 0.001$, partial $\eta^2 > 0.272$), sequence length reduced response time in PRE and REF but not in POST (log(RT): condition × sequence length: $F_{4,144} = 33.1$, $p < 0.001$, partial $\eta^2 = 0.479$; POST: $F_{2,72} = 0.74$, $p = 0.45$, partial $\eta^2 = 0.020$; $BF_{01} = 13.38$; PRE: $F_{2,72} = 37.2$, $p < 0.001$, partial $\eta^2 = 0.508$; REF: $F_{2,72} = 62.5$, $p < 0.001$, partial $\eta^2 = 0.634$; **Supplementary Fig. 2b**).

To better understand this difference between conditions, we regressed the effect of sequence length together with the effect of the strength of evidence provided by each sequence – computed as the log-likelihood ratios, see **Methods** – against response times (**Supplementary Fig. 2c**). We first signed the strength of evidence according to participants’ choices and log-transformed the response times. While the strength of evidence provided by the sequence similarly speeds up the response times in all conditions (coefficient $b_1$: $t$-test against 0: all $t_{36} < -12.5$, all $p < 0.001$; PRE-POST-REF: $F_{2,72} = 1.78$, $p = 0.17$, partial $\eta^2 = 0.047$, $BF_{01} = 5.83$), the length of the sequence has a clear opposite effect on the response times in POST compared to the conditions where the category-mapping cue was available (coefficient $b_2$ PRE-POST-REF: $F_{2,72} = 31.2$, $p < 0.001$, partial $\eta^2 = 0.464$). Indeed, the length of the sequence seems to slow down participants when they have to accumulate information in the stimulus space while it speeds them up when they can use their knowledge of the categories ($t$-test against 0, POST: $t_{36} = 3.72$, $p < 0.001$, Cohen’s $d = 0.61$, 95% CI = [0.011 0.041]; PRE: $t_{36} = -2.44$, $p = 0.0197$, Cohen’s $d = -0.40$, 95% CI = [-0.038 -0.003], $BF_{10} = 2.37$; REF: $t_{36} = -4.16$, $p < 0.001$, Cohen’s $d = -0.68$, 95% CI = [-0.059 -0.020]).

To further investigate what might slow down participants in POST, we regressed out two other components, independent from participants’ reported choices, to predict the response times. The first regressor was again the absolute strength of evidence provided by each sequence, but this time irrespective of participants’ choice. The second regressor was the amount of information provided by the sequence when computing the absolute evidence as log-likelihood ratios with respect to the ‘null’ categories located at the boundaries between the true categories. This second regressor reaches its maximal values for stimuli orientations that are totally uninformative for the final choice, while reaching minimal values for orientations closer to the true category mean. We call ‘true’ projection axis the center of the generative category and ‘null’ projection axis the boundary between categories (see also **Fig. 5**). We found a main effect of condition on how the true absolute evidence strength modulated the response times (left; $F_{2,72} = 9.9$, $p < 0.001$, partial $\eta^2 = 0.217$; **Supplementary Fig. 2d**) but this modulation seemed to be driven by REF as it remains comparable between POST and PRE (post-hoc $t$-

test, difference POST-PRE: $t_{36} = 0.34$, $p = 0.93$, Cohen's $d = 0.057$, 95% CI = [-0.012 +0.015], $BF_{01} = 5.34$). In contrast, irrelevant information has opposite effects between conditions as it slows down participants in POST while slightly speeding them similarly up in the other two conditions (right, $t$-test against 0: POST: $t_{36} = 6.82$, $p < 0.001$, Cohen's $d = 1.12$, 95% CI = [0.023 0.043]; PRE: $t_{36} = -4.69$, $p < 0.001$, Cohen's $d = -0.77$, 95% CI = [-0.027 -0.010]; REF: $t_{36} = -2.10$, $p = 0.043$, Cohen's $d = -0.34$, 95% CI = [-0.020 -0.0003], $BF_{10} = 1.26$, difference PRE-REF: $t_{36} = -1.69$, $p = 0.22$, Cohen's $d = -0.27$, 95% CI = [-0.021 +0.004], $BF_{01} = 1.54$; **Supplementary Fig. 2d**). Together, these findings indicate that accumulating evidence in stimulus space delays the resulting decisions as a function of the number of stimuli in the sequence, but also of the decision-irrelevant information that had to be accumulated until the end of the sequence and discarded after the onset of the category-mapping cue. The same 'null' evidence does not slow down decisions when accumulating evidence in category space (in PRE and REF). Nonlinear attractor models<sup>1-3</sup> can explain how the interaction between decision-relevant and decision-irrelevant information during evidence accumulation can create significant interference – interference that is not predicted by linear (and optimal) evidence accumulation.

### Supplementary Methods

#### Formal equivalence between sensory and category space accumulation models

The proposed Bayesian suboptimal inference model (see **Methods**) accumulates the evidence provided by each stimulus as a log-likelihood ratio $\ell_t$, i.e., as evidence that is already categorized. One might argue that this implementation cannot describe how information is processed when categories are unknown, as in POST. Indeed, accumulating evidence in sensory space instead corresponds to tracking the full posterior over the generative mean after each presented orientation (see **Fig. 1a, top**), which corresponds to tracking the mean orientation of the sequence. We implemented a sensory space accumulation model as follows:

In the normative case, the full posterior is computed following Bayes' rule:

$$p(\mu|\theta_{1:n}) = \frac{p(\theta_{1:n}|\mu)p(\mu)}{p(\theta_{1:n})}.$$

As every orientation from the sequence is independently sampled, the likelihood becomes:

$$p(\theta_{1:n}|\mu) = \prod_{t=1}^n p(\theta_t|\mu),$$

and, assuming a uniform category prior $p(\mu)$ and writing $C = \frac{p(\mu)}{p(\theta_{1:n})}$ for a constant term independent of $\mu$, we have:

$$p(\mu|\theta_{1:n}) = C \prod_{t=1}^n p(\theta_t|\mu).$$

In our task, orientations are sampled from a von Mises distribution with fixed concentration $\kappa$, adjusted for the half-circle symmetry of orientations, such that the likelihood of sampling $\theta_t$ for any given mean $\mu$ is given by:

$$p(\theta_t|\mu) = \frac{e^{\kappa \cos(2(\theta_t - \mu))}}{\pi I_0(\kappa)}.$$

The full posterior distribution over all possible $\mu$ becomes:

$$p(\mu|\theta_{1:n}) = C \prod_{t=1}^n \frac{e^{\kappa \cos(2(\theta_t - \mu))}}{\pi I_0(\kappa)} = \frac{C}{D} \exp(\kappa \sum_{t=1}^n \cos(2(\theta_t - \mu))),$$

where $D = (\pi I_0(\kappa))^n$ is a constant term and $I_0(\cdot)$ is the modified Bessel function of order 0.

The cosine can be expressed by taking the real part of the complex number $z = e^{i2(\theta_t - \mu)}$:

$$\cos(2(\theta_t - \mu)) = \text{Re}(e^{i2(\theta_t - \mu)}) = \text{Re}(e^{i2\theta_t} e^{-i2\mu}) = \text{Re}(z_t \cdot \bar{z}_\mu),$$

with $z_t = e^{i2\theta_t}$ the complex number associated with the orientation $\theta_t$ and $\bar{z}_\mu = e^{-i2\mu}$ the complex conjugate of $z_\mu$.

The posterior probability becomes:

$$p(\mu|\theta_{1:n}) = \frac{C}{D} \exp(\kappa \sum_{t=1}^n \text{Re}(z_t \cdot \bar{z}_\mu)).$$

By linearity of the real-part operator:

$$p(\mu|\theta_{1:n}) = \frac{C}{D} \exp\left(\kappa \text{Re}\left(\sum_{t=1}^n z_t \cdot \bar{z}_\mu\right)\right) = \frac{C}{D} \exp\left(\kappa \text{Re}(Z \cdot \bar{z}_\mu)\right) = \frac{C}{D} \exp(\kappa |Z| \cos(2(\bar{\theta} - \mu))),$$

with $Z$ the resultant of the sequence orientations $Z = \sum_{t=1}^n z_t = |Z|e^{i2\bar{\theta}}$.

Therefore, in our task, the full posterior over $\mu$ depends on the stimulus sequence only through the resultant complex vector $Z$ with norm $|Z|$ and mean orientation $\bar{\theta}$.

At the end of a sequence, the probability of each category $A$ and $B$ is given by evaluating the posterior over the generative mean $\mu$ at $\mu_A$ and $\mu_B$:

$$p(A) = p(\mu = \mu_A | \theta_{1:n}) = \frac{c}{D} \exp(\kappa |Z| \cos(2(\bar{\theta} - \mu_A))),$$

$$p(B) = p(\mu = \mu_B | \theta_{1:n}) = \frac{c}{D} \exp(\kappa |Z| \cos(2(\bar{\theta} - \mu_B))).$$

The sign of the log-posterior ratio provides the most likely category:

$$\mathcal{L}_{\text{sensory}} = \log \frac{p(\mu = \mu_A | \theta_{1:n})}{p(\mu = \mu_B | \theta_{1:n})} = \log \frac{\exp(\kappa |Z| \cos(2(\bar{\theta} - \mu_A)))}{\exp(\kappa |Z| \cos(2(\bar{\theta} - \mu_B)))} = \kappa |Z| (\cos(2(\bar{\theta} - \mu_A)) - \cos(2(\bar{\theta} - \mu_B))).$$

Given that the categories $A$ and $B$ are orthogonal $\mu_B = \mu_A + \pi/2$,

$$\mathcal{L}_{\text{sensory}} = 2\kappa |Z| (\cos(2(\bar{\theta} - \mu_A))).$$

This is proportional to the projection of the resultant vector onto the category-A axis, which is the real part of their complex inner product:

$$\mathcal{L}_{\text{sensory}} = 2\kappa \text{Re}(Z \cdot \overline{z_{\mu_A}}).$$

This is identical to the log-posterior ratio of the category space model:

$$\mathcal{L}_{\text{category}} = \sum_{t=1}^n \ell_t = \sum_{t=1}^n 2\kappa \cos(2(\theta_t - \mu_A)) = 2\kappa \sum_{t=1}^n \text{Re}(z_t \cdot \overline{z_{\mu_A}}) = 2\kappa \text{Re}(\sum_{t=1}^n z_t \cdot \overline{z_{\mu_A}}) = 2\kappa \text{Re}(Z \cdot \overline{z_{\mu_A}}) = \mathcal{L}_{\text{sensory}}.$$

We introduce suboptimality in the sensory space accumulation model similarly to the category space accumulation model:

1. a time-invariant exponential inference imbalance $\gamma_{\text{sensory}}$ capturing the relative gain applied to the previously accumulated resultant vector during the accumulation process:

$$Z = \sum_{t=1}^n z_t \cdot \exp(\gamma_{\text{sensory}})^{n-t}$$

for $\gamma < 0$, previous resultant vectors are down-weighted, similarly to an inference leak, since $\exp(\gamma) < 1$, producing a recency effect; $\gamma > 0$ leads to a primacy effect by putting more weight on early accumulated observations and $\gamma = 0$ corresponds to a Bayes-optimal weighting of prior and likelihood.

2. inference noise $\sigma_{\text{inf sensory}}$ capturing variability at the inference stage through isotropic complex Gaussian perturbations of each stimulus vector resulting in:

$$\text{Re}(\hat{z}_t) \sim \mathcal{N}(\text{Re}(z_t), \sigma_{\text{inf sensory}}^2)$$

$$\text{Im}(\hat{z}_t) \sim \mathcal{N}(\text{Im}(z_t), \sigma_{\text{inf sensory}}^2)$$

Both suboptimality parameters are equivalent to their category-space counterparts: projecting the isotropic noise onto the category axis yields Gaussian noise on each log-likelihood with s.d. $2\kappa\sigma_{\text{inf sensory}}$, which equals $\sigma_{\text{inf sensory}}$ for $\kappa = 0.5$. Noise along the orthogonal dimension is discarded by the final projection, such that the two models make identical predictions. We fitted this sensory-space model to participants' choices (same fitting procedure as the category-space model, see **Methods**) and confirmed the equivalence of the two models by correlating their best-fitting parameters in each condition (sensory versus category inference imbalance $\gamma$: all $r^2 > 0.98$; sensory versus category inference noise $\sigma_{\text{inf}}$: all $r^2 > 0.98$; sensory versus category response lapse $\varepsilon$: $r^2 > 0.96$; **Supplementary Fig. 3**).

### Supplementary References

1. Amari, S. Dynamics of pattern formation in lateral-inhibition type neural fields. *Biol. Cybern.* **27**, 77–87 (1977).
2. Ermentrout, B. Neural networks as spatio-temporal pattern-forming systems. *Rep. Prog. Phys.* **61**, 353–430 (1998).
3. Esnaola-Acebes, J. M., Roxin, A. & Wimmer, K. Flexible integration of continuous sensory evidence in perceptual estimation tasks. *Proc. Natl Acad. Sci. USA* **119**, e2214441119 (2022).
